# Rho-ROCK signaling and α-Catenin mediate β-Catenin-driven hyperplasia in the adrenal via adherens junctions

**DOI:** 10.1101/2025.10.06.680572

**Authors:** Mesut Berber, Betul Haykir, Nick A. Guagliardo, Vasileios Chortis, Kleiton Silva Borges, Paula Q. Barrett, Felix Beuschlein, Diana L. Carlone, David T. Breault

## Abstract

How β-Catenin (βCat) mediates tissue hyperplasia is poorly understood. To explore this, we employed the adrenal cortex as a model system given its stereotypical spatial organization and the important role βCat plays in homeostasis and disease. For example, excessive production of aldosterone by the adrenal cortex (primary aldosteronism, PA) constitutes a significant cause of cardiovascular morbidity, which has been associated with βCat gain-of-function (βCat-GOF). Adherens junctions (AJs) connect the actin cytoskeletons of adjacent zona Glomerulosa (zG) cells via a cadherin/βCat/α-Catenin (αCat) complex and mediate aldosterone production. Whether βCat-GOF drives zG hyperplasia, a key feature of PA, via AJs is unknown. Here, we show that aldosterone secretagogues (K^+^, AngII) and βCat-GOF mediate AJ enrichment via Rho-ROCK-actomyosin signaling. In addition, Rho-ROCK inhibition leads to altered zG rosette morphology and decreased aldosterone production. Mice with zG-specific βCat-GOF demonstrate increased AJ formation and zG hyperplasia, which was blunted by Rho-ROCK inhibition and deletion of αCat. Further, analysis of human aldosterone-producing adenomas (APAs) revealed high levels of βCat expression were associated with increased membranous expression of K-Cadherin. Together, our findings identify Rho-ROCK signaling and αCat as key mediators of AJ enrichment and β-Catenin-driven hyperplasia.

**One Sentence Summary:** This study demonstrates that β-Catenin-driven hyperplasia in the adrenal cortex, a key feature of primary aldosteronism, is mediated through Rho-ROCK signaling and α-Catenin-dependent stabilization of adherens junctions, with significant implications for patients with primary aldosteronism.

**Highlights:**

- Rho-ROCK signaling drives AJ enrichment in the adrenal
- ROCK inhibition via fasudil blunts aldosterone production
- βCat drives adrenal hyperplasia via enhanced AJ enrichment
- ROCK inhibition or ɑCat deletion block zG hyperplasia

## INTRODUCTION

Primary aldosteronism (PA) is caused by aldosterone-producing micronodules (APMs) and larger adenomas (APAs) arising from zona glomerulosa (zG) cells of the adrenal cortex, which are characterized by autonomous over-production of aldosterone (*1*). PA represents the most common cause of secondary hypertension and includes 25% of patients with resistant hypertension (*2*). Delays in the diagnosis and treatment of PA result in a higher risk of cardiovascular morbidity including stroke, heart failure, renal damage and overall morbidity (*3*). While the majority of APMs and APAs arise from somatic mutations in ion channels associated with aldosterone production (*1*), the underlying cellular mechanisms driving hyperplasia within these lesions remain incompletely understood(*4*). Moreover, it is unclear to what extent these mechanisms intersect with pathways activated by plasma potassium (K^+^) or angiotensin II (AngII) signaling–the principal physiological regulators of zG size and aldosterone production (*5*).

Adherens junctions (AJs) are crucial mediators of cell-cell adhesion, structurally linking the actomyosin cytoskeleton of neighboring cells, and play a critical role in epithelial morphogenesis and tissue remodeling (*6*). AJs are composed of core structural elements, including transmembrane cadherins, β-Catenin (βCat), and α-Catenin (αCat). Among the core AJ components, αCat functions as an essential linker protein connecting the cadherin-βCat complex to the actomyosin cytoskeleton of the cell (*7*). AJs orchestrate rosette formation—a conserved developmental mechanism driving organogenesis and epithelial remodeling (*8*). Rosettes arise when five or more epithelial cells reorganize their cell-cell junctions into a flower-like structure. This process occurs through AJ dynamics and actomyosin-driven cytoskeletal remodeling, facilitating organogenesis in systems ranging from *Drosophila* epithelia to the adrenal zG (*9, 10*). In the adrenal, the zG consists of Laminin β1-encapsulated glomerular structures composed of multicellular rosettes (*10*), and in humans, expanded rosettes have been identified in aldosterone-producing cell clusters (*11*), a possible precursor to aldosterone-producing adenomas (APAs).

AJ enrichment results from the dynamic balance between AJ assembly and disassembly, a process that is tightly regulated to allow cells to modulate their adhesion strength and respond to physiological cues during tissue homeostasis (*12*). Thus, dysregulation of AJ dynamics can significantly impact tissue structure and function. For example, enhanced AJs are associated with the initiation of tumor formation, while weakened AJs are associated with epithelial-mesenchymal transition, which can ultimately result in tumor cell metastasis (*13–15*). Morphologically, AJs exist in two distinct forms: punctate AJ (pAJ) and zonula adherens (ZA) (*6, 16, 17*). pAJs are associated with transient weakening of AJs, enabling cell movement and tissue rearrangement, and consist of discrete points of cell adhesion. In contrast, ZAs demonstrate increased strength and stability, are essential for the maintenance of tissue integrity, and are composed of continuous and dense AJ clusters that span cell-cell boundaries. Various signaling pathways have been implicated in the regulation of AJ stability, including those downstream of Rho-GTPases (*9, 18*). For example, Rho-associated, coiled coil-containing kinase (ROCK) increases non-muscle Myosin II (NMII) motor activity through phosphorylation of the regulatory light chain of NMII (*18*). Moreover, the recruitment of NMII to the AJ and the motor activity of NMII are critical for the formation, expansion, and maintenance of ZAs (*19*). Thus, elucidating pathways that regulate AJs in the zG could constitute a significant advance in understanding the cellular mechanisms underlying zG hyperplasia.

While βCat functions as an AJ linker protein, it also serves as an essential transcription factor in the canonical WNT/βCat pathway, which is regulated, in part, via continuous degradation (*20*). Stabilization of βCat, via phosphorylation or deletion of key degron domains, has been implicated in zG pathology in both humans and mice (*21, 22*). For example, somatic gain-of-function (GOF) mutations in *CTNNB1* (the gene encoding βCat) (βCat-GOF) result in increased aldosterone production (*23, 24*), and βCat stabilization is detected in upwards of 70% of APAs (*24*). Moreover, zG-specific βCat-GOF in mice leads to zG hyperplasia through a block in transdifferentiation of zG cells into zona Fasciculata cells without a corresponding increase in proliferation (*22, 24, 25*). Despite the central importance of βCat-GOF in zG hyperplasia and APA formation, it remains unclear whether these effects are mediated through altered gene expression, regulation of AJs or both.

Here, we investigate the role of Rho-ROCK signaling, αCat and βCat-GOF as mediators of AJ stability and zG hyperplasia. We discovered that physiological stimulation with K^+^ enhanced AJ stability in the zG via Rho-ROCK signaling. In addition, βCat-GOF in the zG led to AJ enrichment, and disruption of AJs (via inhibition of Rho-ROCK signaling), blocked βCat-driven zG hyperplasia, underscoring a key role for AJ stability as a driver of zG hyperplasia. Consistent with this, deletion of αCat disrupted zG morphology and blocked βCat-driven zG hyperplasia. Finally, in human APAs, levels of βCat positively correlated with levels of K-Cadherin (KCad), consistent with βCat being a driver of stability, which has important implications for the development of targeted therapies for patients with PA.

## RESULTS

### K^+^ and AngII stimulation enhance AJ stability via Rho-ROCK signaling

To identify physiological mechanisms regulating zG function, we reanalyzed our phosphoproteomic dataset examining the effects of K^+^ stimulation on aldosterone production in the human NCI-H295R adrenocortical cell line (*26*). Gene Ontology and Reactome Pathway analyses revealed that K^+^ stimulation led to increased levels of phosphoproteins involved in cadherin binding, actin binding, and Rho-GTPase signaling, each of which is associated with AJ enrichment (**Supplementary Fig. 1A**). To extend these findings, we assessed the effects of the aldosterone secretagogues K^+^ and AngII on AJs (including the transition from pAJs to ZA) (**Fig. 1A, Supplementary Fig. 1B**). First, we used co-immunoprecipitation (co-IP) for αCat in NCI-H295R cells treated with either K^+^ or AngII. Both AngII and K^+^ significantly increased KCad association with αCat, suggesting increased AJ stability (**Fig. 1B-C**). To better characterize the K^+^-induced transition from pAJs to more stable ZAs, we developed a quantitative line profile analysis (qLPA), by adapting the fluorescence intensity line profiling techniques in ImageJ into a standardized quantitative approach, where the fluorescence intensity of KCad and αCat was measured along 50-pixel-wide lines drawn between adjacent nuclei and normalized to a 0-100 point scale. This method allowed for direct comparison of AJ formation across experimental conditions. Consistent with the co-IP results, qLPA following stimulation with K^+^ revealed an increase in the transition from pAJs to ZAs, and increased AJ formation (**Fig. 1D-F, Supplementary Fig. 1C, 2A**). Similar results were observed following stimulation with AngII (**Supplementary Fig. 1D, 2E**). Considering that both K^+^ and AngII converge on Ca^2+^ signaling, we inhibited L-type Ca^2+^ channels with nifedipine to determine the role of Ca^2+^ influx in AJ assembly. As predicted, nifedipine prevented K^+^-induced AJ enrichment **Fig. 1D-F, Supplementary Fig. 2A)**. Together, these data suggest that aldosterone secretagogues led to AJ enrichment through Ca^2+^-mediated signaling between zG cells.

**Figure 1.**
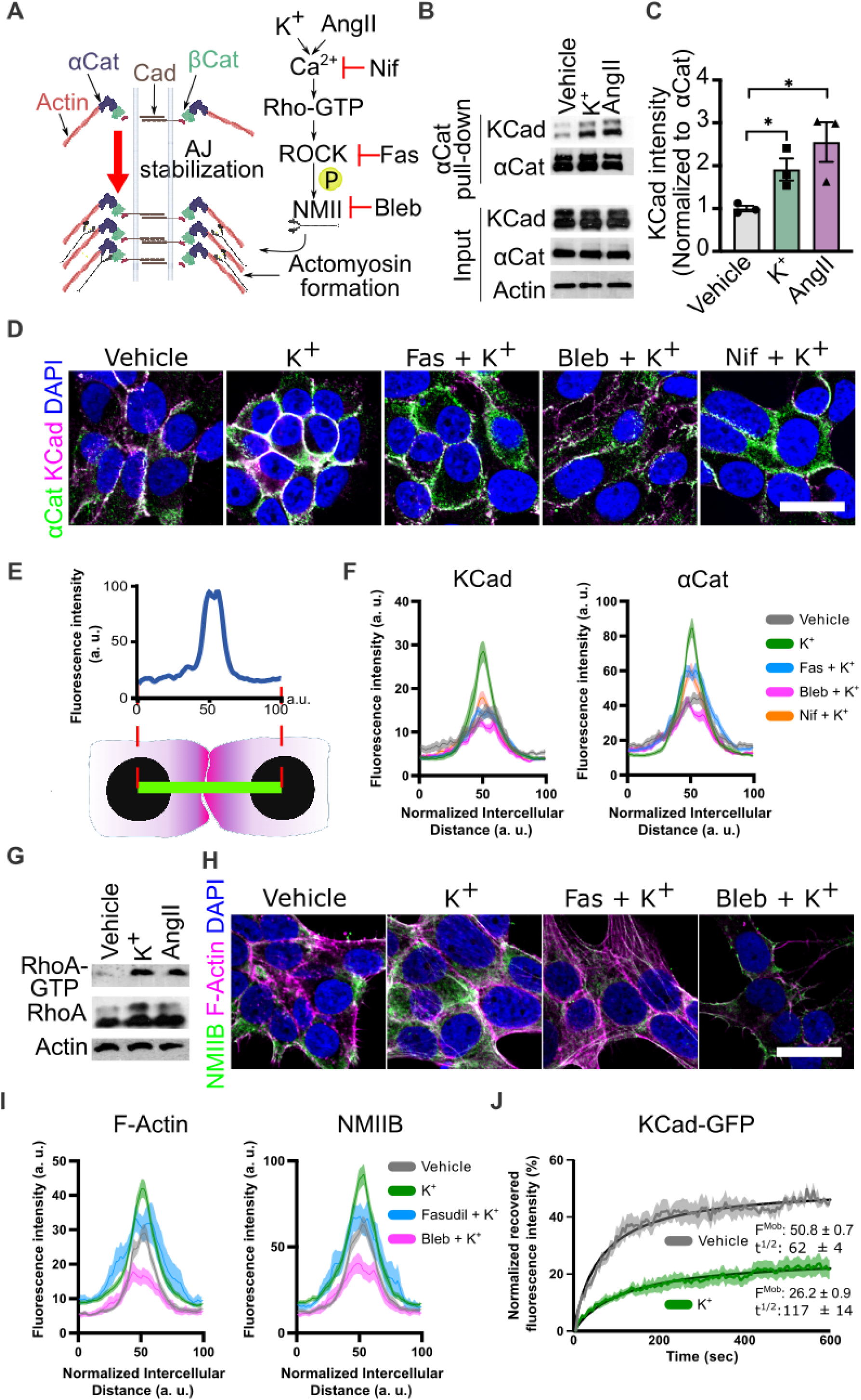
Aldosterone secretagogues increase adherens junction stability via Rho-ROCK-NMII pathway. **A)** Schematic illustrating Rho-ROCK-NMII pathway regulation of adherens junction (AJ) stability. pAJ, puncta AJ; ZA, zonula adherens. **B)** α-Catenin (αCat) co-immunoprecipitation (co-IP) in NCI-H295R cells treated with vehicle (11 mM NaCl), potassium (K^+^; 11 mM KCl) or angiotensin II (AngII; 100 nM) for 48 h. Immunoblots of K-Cadherin (KCad) and αCat are shown. Input, whole cell lysates. **C)** Densitometric quantification of KCad/αCat co-IP (n=3 independent experiments). **D)** Representative image of αCat and KCad immunofluorescence in NCI-H295R cells stimulated with vehicle, K^+^ ± fasudil (10 µM), blebbistatin (10 µM) or nifedipine (10 µM) for 48 h (1 h preincubation with inhibitors). **E)** Schematic of line profile analysis methodology. **F)** Quantitative line profile analysis (qLPA) of αCat and KCad fluorescence intensities represented in (D) (n=24-47 cell-cell interfaces). Lines represent mean and shaded areas represent standard error of the mean (SEM). **G)** RhoA activity in NCI-H295R cells treated with vehicle, K^+^ or AngII for three hours was assessed by pull-down assay using Rhotekin-RBD protein beads. Immunoblots show active RhoA in precipitates (top panel), total RhoA in cell lysates (middle panel), and actin in cell lysates (bottom panel) as a loading control. **H)** Representative images of non-muscle myosin IIB (NMIIB) immunofluorescence and filamentous actin (F-actin) staining in NCI-H295R cells treated as in (D). **I)** qLPA of NMIIB and F-actin fluorescence intensities represented in (H) (n=16-42 cell-cell interfaces). **J.** Normalized KCad-GFP fluorescence recovery after photobleaching (FRAP) in NCI-H295R cells treated with vehicle (11 mM NaCl) or potassium (11 mM KCl) for 48 hours. Lines represent mean and shaded areas represent SEM. Statistical significance determined by unpaired two-tailed Student’s *t*-test (*P < 0.05). Data are presented as mean ± SEM. Nuclei were counterstained with DAPI and are shown in blue. Scale bars, 20 μm.

Our phosphoproteomic analysis also identified Rho GTPase signaling as a putative downstream mediator of K^+^ stimulation (**Supplementary Fig. 1A**), indicating this pathway may mediate the enhancement of AJ stability. To confirm this, we treated NCI-H295R cells with K^+^ or AngII, which revealed increased RhoA-GTP activation (**Fig. 1G**). Next, we assessed actomyosin formation, a downstream marker of RhoA activity (**Fig. 1A**) (*27*), by staining for myosin light chain 2 (MLC2), filamentous actin (F-Actin) and non-muscle myosin IIb (NMIIB). K^+^ stimulation promoted MLC2 phosphorylation and robust actomyosin assembly at cell-cell contacts, which directly correlated with increased fluorescence intensity and more continuous distribution of KCad and αCat along the normalized membrane interface, reflecting the molecular basis for the observed transition from pAJs to more stable ZAs, as assessed by qLPA (**Fig. 1H-I, Supplementary Fig. 2B-C**). Consistent with these findings, inhibition of Rho-associated protein kinase (ROCK) activity (with fasudil) (*28, 29*) or myosin motor activity (with blebbistatin) (*30*) abrogated ZAs and actomyosin formation (**Fig. 1H-I**). Indeed, inhibition of ROCK and NMII activity using fasudil, Y27632, and blebbistatin disrupted the membrane localization and junction formation of KCad and αCat, as evidenced by reduced fluorescence intensity at cell-cell interfaces (**Fig. 1D, F, Supplementary Fig. 2**). Similar results were observed following stimulation with AngII (**Supplementary Fig. 1D, 2E**). Finally, to establish whether increased AJ stability contributes to the increase in AJ enrichment seen following K^+^ stimulation, we performed fluorescence recovery after photobleaching (FRAP) using a KCad-GFP fusion protein in H295R cells. The results demonstrated that K^+^ stimulation significantly decreased the mobile fraction of KCad (thereby increasing the stable fraction) indicating K^+^ stimulation led to increased AJ stability **(Fig 1J)**. Together, these data indicate that K^+^ and AngII promote AJ enrichment and stability via Rho-ROCK-NMII signaling in adrenocortical cells.

### Rho-ROCK signaling mediates zG morphology and aldosterone production

Given AJs are crucial for normal zG morphology (i.e., zG rosettes), we next assessed whether Rho-ROCK signaling mediated this process. We treated two-month-old wild-type mice with fasudil for four weeks and assessed zG structure, gene expression and aldosterone production. Compared to vehicle-treated control mice, fasudil treatment led to a significant reduction in 2D glomerular area (a measure of zG morphology assessed using immunostaining for Laminin β1) and zG rosettes, despite no change in the overall size of the zG (assessed using immunostaining for Dab2) (**Fig. 2A-D**). Next we assessed the functional impact on the adrenal of reduced 2D glomerular area and rosettes in fasudil-treated mice, which revealed a decrease in *Cyp11b2* (encoding aldosterone synthase) expression (**Fig. 2E**), and reduced aldosterone levels (**Fig. 2F**), with a trend toward increased renin expression levels (**Fig. 2G**). To confirm the reduction in aldosterone levels, we repeated the analysis with a more rigorous two-week fasudil treatment protocol in control mice, incorporating pre- and post-treatment aldosterone measurements, which verified decreased aldosterone production following fasudil treatment (**Fig 2H**). Taken together, these results indicate a direct impact of fasudil on adrenal function and rosette maintenance, the modest changes in renal renin mRNA levels may reflect the complex, multi-level regulation of renin expression and suggest that both direct adrenal effects and compensatory renal responses may contribute to the observed hormonal changes in this well-coordinated endocrine system.

**Figure 2.**
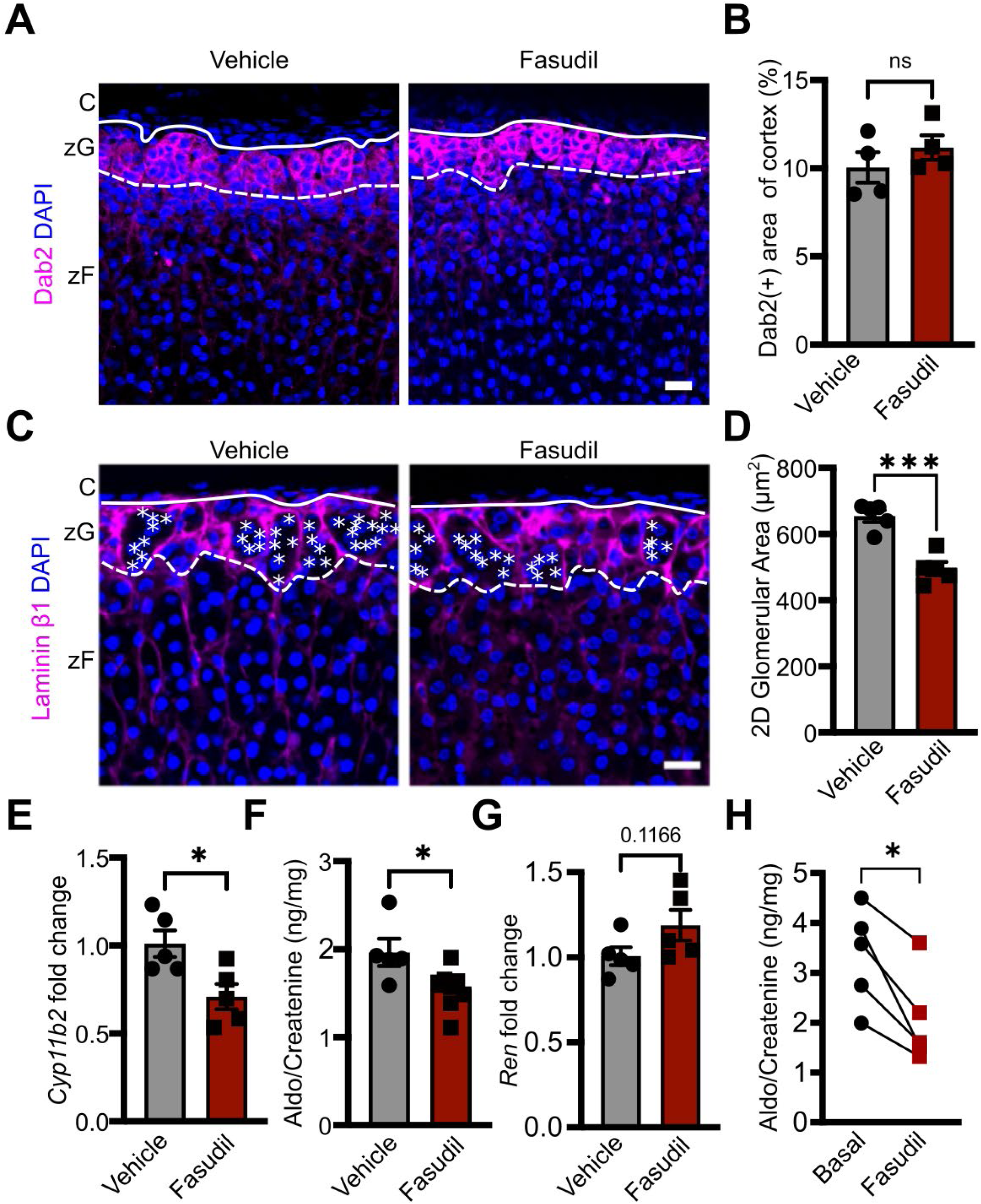
ROCK inhibition via fasudil decreases 2D glomerular area in the zG and blunts aldosterone production. **A)** Representative images of Dab2 immunofluorescence in adrenal sections from two-month-old control female mice treated with either vehicle (PBS) or fasudil (30 mg/kg) administered every other day for 28 days. **B)** Quantification of the percentage of Dab2(+) area in the adrenal cortex, as shown in (A) (n = 4, 4 mice). **C)** Laminin β1 immunofluorescence in adrenal sections from two-month-old female C57BL/6 mice treated as in (A). Images highlight the glomerular structures within the zona glomerulosa (zG) region of the adrenal cortex. Solid lines mark the boundary between the adrenal capsule (C) and zona glomerulosa (zG), while dashed lines indicate the boundary between the zG and zona fasciculata (zF). Each white asterisk denotes an individual cell within a zG rosette. **D)** Quantification of two-dimensional (2D) glomerular area via Laminin β1 labeling represented in (C) (n = 5, 5 mice). **E)** *Cyp11b2* mRNA expression in adrenals of vehicle- and fasudil-treated two-month-old female C57BL/6 mice, assessed by quantitative PCR (qPCR) (n = 5, 5 mice). **F)** Measurement of 24-hour urinary aldosterone (Aldo) levels in two-month-old female C57BL/6 mice treated as in (A), assessed by radioimmunoassay and normalized to creatinine (n = 5, 8 mice). **G)** *Renin* mRNA expression in kidneys of two-month-old female C57BL/6 mice treated as in (A), assessed by qPCR (n = 5, 5 mice). **H)** Measurement of 24-hour urinary aldosterone levels in 7-12-month-old control male mice before (basal) and after fasudil treatment (30 mg/kg) administered 6 times weekly for 14 days, assessed by RIA and normalized to creatinine (n = 5 mice). Statistical significance determined by unpaired two-tailed Student’s *t*-test or ratio paired t-test (in H) (*P < 0.05, ***P <0.001, ns, not significant). Data are presented as mean ± SEM. Scale bars, 20 μm.

### βCat stabilization promotes AJ stabilty

While βCat is a core component of the AJ complex, it is not known whether increased levels of βCat expression might directly affect AJ structures. To explore this, we assessed the impact of βCat stabilization on AJ integrity. First, we treated NCI-H295R cells with CHIR 99021 (CHIR), an inhibitor of GSK3β, to enhance βCat stabilization (**Supplementary Fig. 3A-C**). Remarkably, treatment with CHIR significantly enhanced membrane localization of KCad, αCat, and F-actin, leading to the formation of well-defined ZAs at cell-cell contacts, compared with pAJs in vehicle-treated cells (**Fig. 3A-C**). Moreover, treatment with CHIR led to a pronounced aggregation of NCI-H295R cells into rosette-like clusters, reminiscent of the spatial organization observed in zG rosettes (**Supplementary Fig. 3D**). Next, we assessed the impact of inhibiting the Rho-ROCK-NMII signaling pathway (using fasudil or blebbistatin) on CHIR-treated cells, which revealed markedly reduced AJ enrichment and attenuation of ZAs formation (**Fig. 3A-C, Supplementary Fig. 3E-F**). Similar results were observed following treatment with the ROCK inhibitor Y27632 (**Supplementary Fig. 3B**). These findings demonstrate that the Rho-ROCK signaling axis is essential for βCat-mediated enhancement of AJ enrichment and multicellular organization in adrenocortical cells, providing mechanistic insight into how tissue architecture may be regulated in the zG.

**Figure 3.**
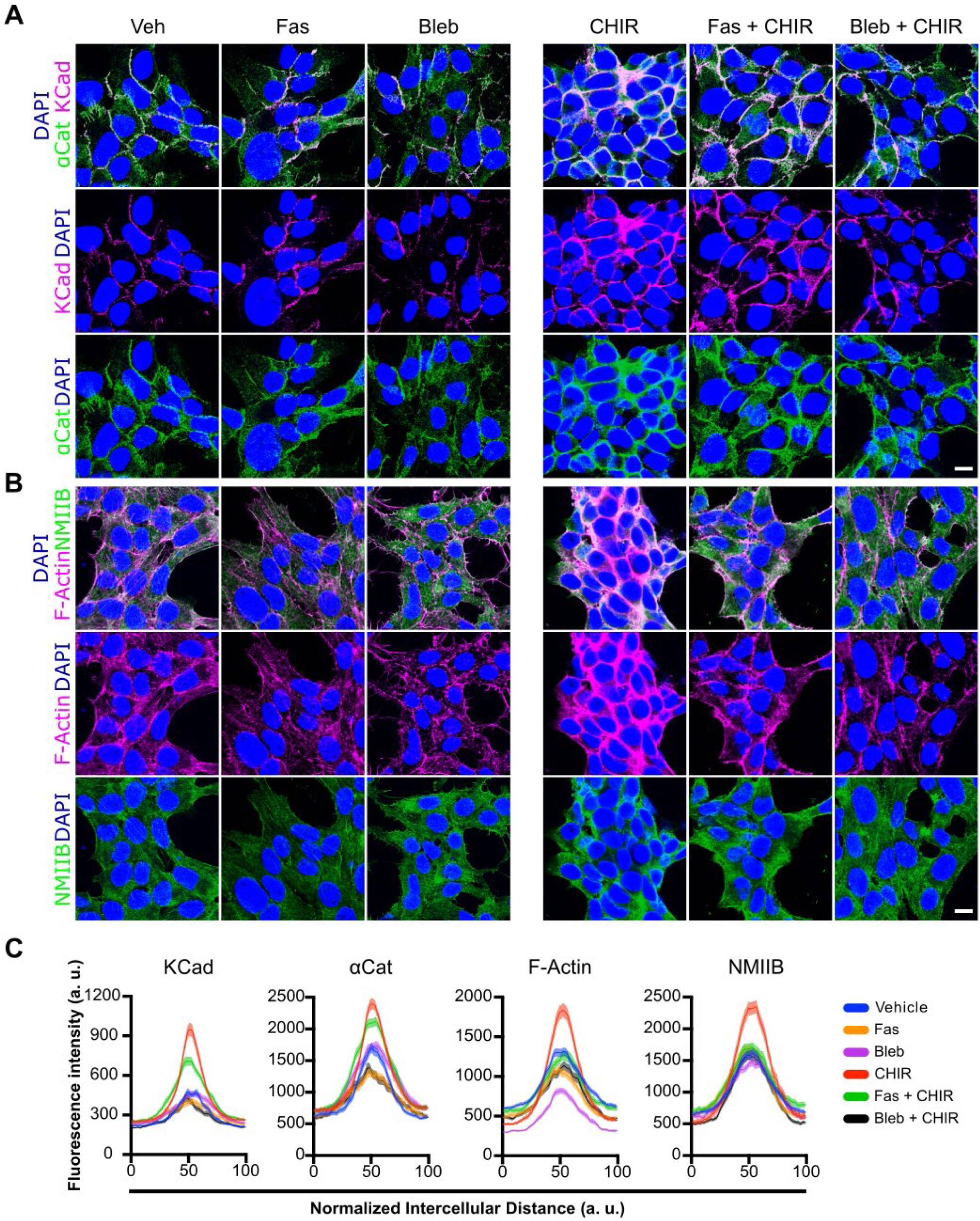
β-Catenin stabilization via CHIR stimulation enhances adherens junction stability. **A)** Representative images of α-Catenin (αCat) and K-Cadherin (KCad) immunofluorescence in NCI-H295R cells stimulated with Vehicle (Veh; DMSO), CHIR 99021 (CHIR; 5µM) ± fasudil (10 µM) or blebbistatin (10 µM) for 48 h (1 h preincubation with inhibitors). **B)** Representative images of Non-muscle myosin IIB (NMIIB) immunofluorescence and filamentous actin (F-actin) staining in NCI-H295R cells treated as in (A). **C)** Quantitative line profile analysis of KCad, αCat, F-actin, and NMIIB fluorescence intensities from (A, B) (n=64-121 cell-cell interface). Lines represent mean and shaded areas represent standard error of the mean (SEM). Scale bars, 20 µM.

We previously showed that βCat stabilization drives progressive zG hyperplasia and increases zG-rosette structures, *in vivo* (*10, 22*). To assess whether these effects of βCat stabilization are mediated through AJ enrichment, we employed zG-specific *Cyp11b2^Cre/wt^::Ctnnb1^flox(Ex3)/wt^* mice, referred to as βCat-GOF (*22*) and *Cyp11b2^Cre/wt^::Ctnnb1^wt/wt^* mice (*31*), which were used as controls. Analysis of βCat-GOF and control mice confirmed zG hyperplasia, as evidenced by an increased number of Dab2(+) cells (**Supplementary Fig. 4A**). Since βCat functions as a transcription factor in addition to its role in the AJ, we first assessed whole-transcriptome data from βCat-GOF and control adrenals (*10*). This analysis revealed increased expression of genes encoding core AJ components, including *Ctnna1* (αCat), *Ctnnd1* (p120-catenin, p120), and *Cdh6* (K-Cadherin, KCad) (**Supplementary Fig. 4B-C**) (*32*). The binding of p120 to KCad serves as an established marker of AJ stability (*33*). Next, to assess changes in AJ stability (i.e., conversion of pAJs to ZAs) we performed immunostaining for KCad and p120 in the zG of βCat-GOF and control mice. In line with our previous findings (*10*), we observed a predominance of pAJs in the control zG and a marked increase in ZAs in the βCat-GOF zG (**Fig. 4A & Supplementary Fig. 4C**). To quantify AJ stability, we employed proximity ligation assay (PLA; a sensitive method for visualizing protein-protein interaction *in situ* (*34*)) to target KCad and p120, which revealed a marked increase in the interaction of these AJ proteins in the zG of βCat-GOF mice compared with controls (**Fig. 4B-C, Supplementary Fig. 4D**). Together, these data indicate that βCat stabilization led to increased expression of AJ components as well as AJ stability.

**Figure 4.**
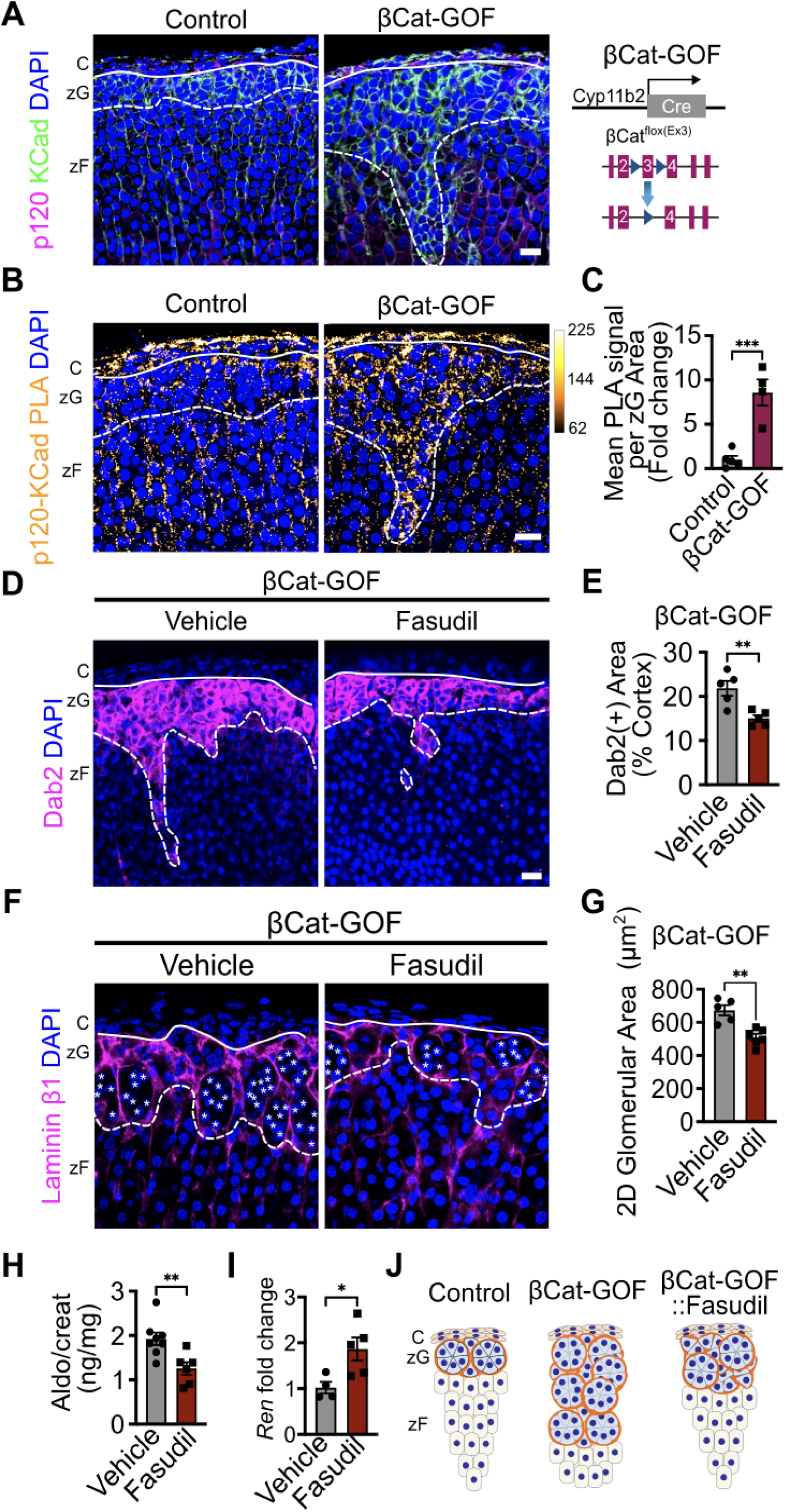
β-Catenin stabilization strengthens adherens junctions in the zona glomerulosa, while ROCK inhibition with fasudil prevents hyperplasia and reduces aldosterone production. **A)** Representative images of K-Cadherin (KCad) and p120-Catenin (p120) immunofluorescence in adrenal sections from two-month-old control and βCat-GOF female mice. Schematic of Cyp11b2^Cre/+^::Ctnnb1^flox(Ex3)/wt^ (βCat-GOF) mice. **B)** Representative images of KCad/p120 proximity ligation assay (PLA) signal in adrenal sections from two-month-old control and βCat-GOF female mice. **C)** Quantification of PLA signal per zona glomerulosa (zG) area from control and βCat-GOF mice represented in (B) (n = 5, 4 mice). **D)** Representative images of Dab2 immunofluorescence in adrenal sections from two-month-old βCat-GOF female mice treated with either vehicle (PBS) or fasudil (30 mg/kg) administered every other day for 28 days. **E)** Quantification of the percentage of Dab2(+) area in the adrenal cortex, as shown in (D) (n = 5, 5 mice). **F)** Representative images of Laminin β1 immunofluorescence in adrenal sections from βCat-GOF female mice treated as in (D). Each white asterisk denotes an individual cell within a zG rosette. **G)** Quantification of the two-dimensional (2D) area of glomerular structures, as defined by Laminin β1 labeling in (F) (n = 5, 6 mice). **H)** Measurement of 24-hour urinary aldosterone (Aldo) levels of four-month-old βCat-GOF female mice treated with either vehicle (PBS) or fasudil (30 mg/kg) administered six times per week for 28 days, assessed by radioimmunoassay and normalized to creatinine (Creat) (n = 8, 6 mice). **I)** *Renin* (*Ren*) mRNA expression in kidneys of four-month-old βCat-GOF female mice treated as in (H), assessed by quantitative PCR (n = 4, 5 mice). **J)** Schematic of control, βCat-GOF and βCat-GOF::fasudil-treated mice. All statistical significance determined by unpaired two-tailed Student’s *t*-test (*P < 0.05, **P < 0.01, ***P < 0.001). Data are presented as mean ± SEM. Solid lines mark the boundary between the adrenal capsule (C) and zona glomerulosa (zG), while dashed lines indicate the boundary between the zG and zona fasciculata (zF). Scale bars, 20 µM.

To evaluate the functional relationship between Rho-ROCK signaling and βCat-driven zG hyperplasia, we administered fasudil or vehicle to βCat-GOF mice at two months of age in female mice. Fasudil treatment of βCat-GOF mice led to a marked reduction in Dab2(+) zG area (**Fig. 4D-E**). In addition, fasudil treatment resulted in a significant reduction in zG glomerular area and zG rosettes (**Fig. 4F-G**). Next, given the progressive nature of the βCat-GOF phenotype with age (*22*), we confirmed these findings in both male and female mice at four months of age, (**Supplementary Fig. 5A-D**). In addition, we showed treatment of βCat-GOF mice (with established zG cell hyperplasia) with fasudil led to decreased aldosterone production with a corresponding increase in renin expression (**Fig. 4H-I, Supplementary Fig. 5E-F**) In aged βCat-GOF mice(*22*), which display pronounced aldosterone elevation, two weeks of fasudil treatment reduced 24h urinary aldosterone excretion to levels below those of littermate controls. **(Supplementary Fig. 5G).** Together, these findings establish that stabilization of βCat led to AJ enrichment and disruption of AJs through inhibition of Rho-ROCK signaling blunted βCat-GOF-driven hyperplasia and aldosterone production (**Fig. 4J**).

### zG-specific αCat deletion prevents βCat-GOF-driven zG hyperplasia

Because βCat functions as both a transcription factor and a core AJ component, it is challenging to delineate the contribution of these two activities on zG hyperplasia in zG-specific βCat-GOF mice. Indeed, βCat-GOF induced expression of AJ components themselves (**Supplementary Fig. 4B**). However, fasudil treatment, while blocking zG hyperplasia, did not affect expression of βCat’s transcriptional targets **(Supplementary Fig 5H)**, and inhibiting βCat’s transcriptional activity with ICRT14 failed to prevent CHIR-induced KCad enrichment at cell-cell junctions in H295R cells (**Supplementary Figs. 3A, 6A-B**), suggesting that βcat’s role in AJ may be critical for zG hyperplasia independent of its transcriptional activity. To address this fundamental question and directly assess the role of the AJ in βCat-driven zG hyperplasia and rosette formation, we next targeted αCat, an obligatory AJ adaptor protein that connects the cadherin-catenin complex to the actin cytoskeleton (**Fig. 1A**) (*7*). We first generated zG-specific *Cyp11b2^Cre/wt^::Ctnna1^flox/flox^* mice, referred to as αCat-LOF (loss-of-function) mice. In addition, to directly assess the role of αCat, and by extension AJs, in the context of βCat-driven zG hyperplasia we generated zG-specific *Cyp11b2^Cre/wt^::Ctnnb1^flox(Ex3)/wt^::Ctnna1^flox/flox^* mice, referred to as βCat-GOF::αCat-LOF mice.

Analysis of two-month-old αCat-LOF mice confirmed deletion of αCat in the zG compared with controls (**Supplementary Fig. 6C**). We next assessed the impact of αCat-LOF on zG morphology, which revealed no change in the overall size of the zG compared to controls (**Supplementary Fig. 6D-G**). Despite this, αCat-LOF led to a reduction in zG 2D glomerular area and zG rosette number compared with controls (**Supplementary Fig. 6H-J**). These data indicate that despite no change in overall zG size between αCat-LOF and control mice, αCat is required for normal zG morphology, including formation of rosette structures.

Next, to assess whether loss of αCat alters βCat-driven zG hyperplasia, we compared 2D glomerular area and zG rosette number in control, βCat-GOF, αCat-LOF and βCat-GOF::αCat-LOF mice. As expected, analysis of βCat-GOF mice revealed a 3-fold increase in Dab2(+) zG area (66%±12% of the cortex), compared to control (22%±3%) and αCat-LOF (19%±3%) mice. Remarkably, analysis of βCat-GOF::αCat-LOF revealed a 50% decrease in Dab2(+) zG area (34%±7% of the cortex) compared with βCat-GOF mice (**Fig. 5A-B, Supplementary Fig. 7**). Consistent with these results, we observed similar changes in 2D glomerular area and zG rosette number (**Fig. 5C-E**). These data indicate that αCat-dependent AJ formation is essential for βCat-driven zG hyperplasia (**Fig 5F**).

**Figure 5.**
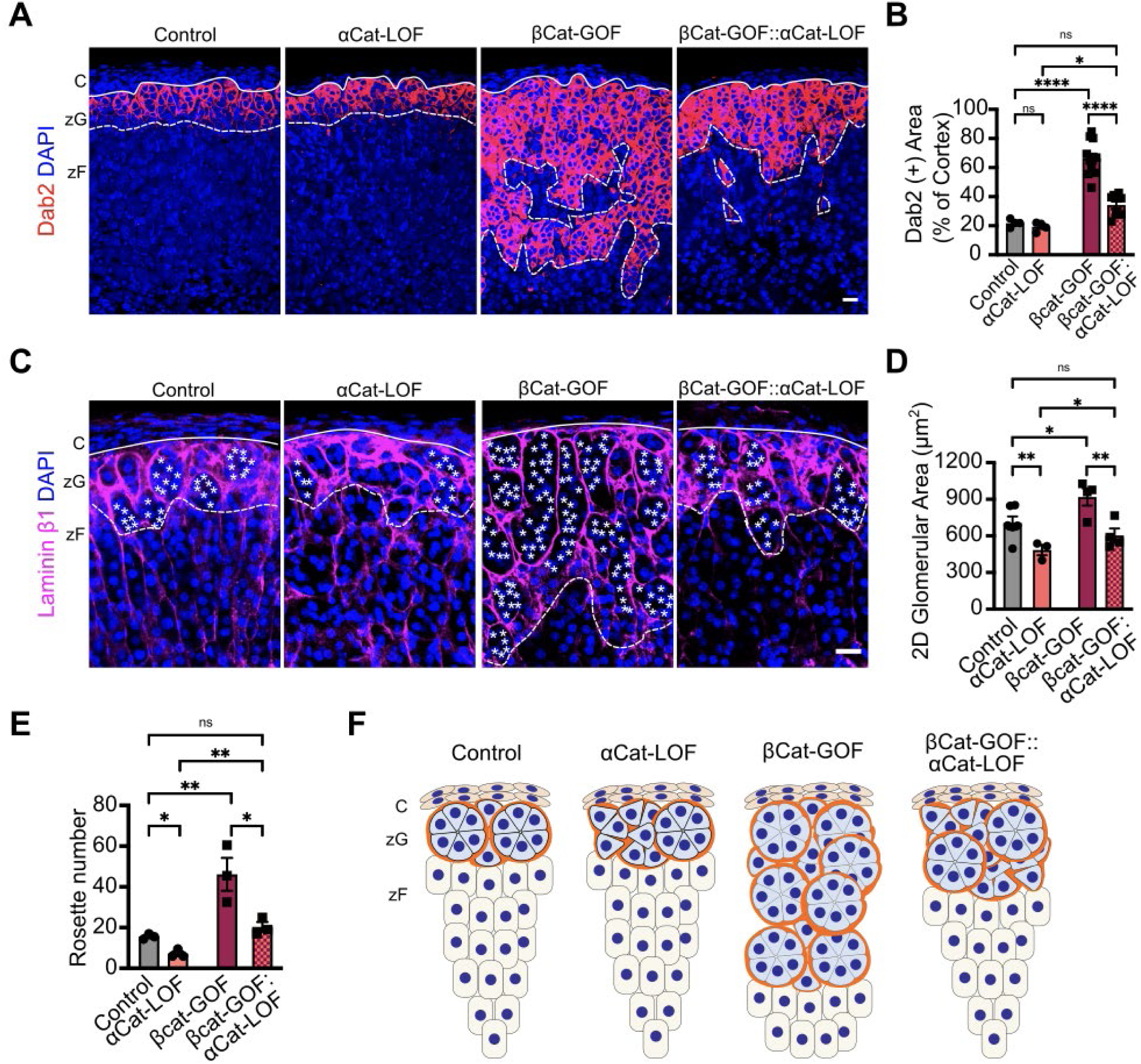
zG-specific α-Catenin deletion attenuates βCat-GOF-induced zG hyperplasia and rosette expansion. **A)** Representative images of Dab2 immunofluorescence in adrenal sections from four-month-old control, αCat-LOF, βCat-GOF, and βCat-GOF::αCat-LOF male mice. **B)** Quantification of the percentage of Dab2(+) area in the adrenal cortex, represented in (A) (n= 4-12 mice). **C)** Representative images of Laminin β1 (Lamb1) immunofluorescence in adrenal sections from four-month-old control, βCat-GOF, αCat-LOF, and βCat-GOF::αCat-LOF male mice. Each white asterisk denotes an individual cell within a zG rosette. **D)** Quantification of the two-dimensional (2D) area of glomerular structures, as defined by Laminin β1 labeling in (C) (n = 3-6 mice). **E)** Quantification of rosette number, defined as clusters of five or more cells within a single glomerular structure measured in regions of 500 x 500 μm, as shown in (C) (n = 3 mice per group). All statistical significance determined by two-way ANOVA with Tukey’s multiple-comparison posttest. *P < 0.05, **P < 0.01, **** < 0.0001, ns, not significant. Data are presented as mean ± SEM. Solid lines mark the boundary between the adrenal capsule (C) and zona glomerulosa (zG), while dashed lines indicate the boundary between the zG and zona fasciculata (zF). Scale bars, 20 µM.

To better understand the mechanism(s) responsible for the rescue of βCat-driven zG hyperplasia with fasudil and loss of αCat, we assessed cell proliferation (immunostaining for Ki67) and cell death (immunostaining for Cleaved Casp3) in the adrenals of βCat-GOF and βCat-GOF::αCat-LOF mice and in βCat-GOF mice treated with fasudil or vehicle. While the number of Ki67(+) cells was comparable between genotypes and treatments, the number of cleaved Casp3(+) cells in the zG was significantly increased in βCat-GOF::αCat-LOF compared to βCat-GOF adrenals and in fasudil-treated βCat-GOF adrenals compared to controls (**Supplementary Fig. 8A-H**). Consistent with these *in vivo* findings, high-dose fasudil induced apoptosis in H295R cells (**Supplementary Fig. 8I**). Together, these results indicate that disruption of AJs—either through αCat-LOF or ROCK inhibition—sensitizes zG cells to apoptosis (*35–37*). Moreover, these findings begin to decouple βCat’s nuclear and AJ functions, revealing AJs as an important factor controlling zG morphology and the hyperplastic zG expansion following βCat-stabilization.

### βCat Expression Correlates with Adherens Junction Enrichment in Aldosterone-Producing Adenomas

Stabilization of βCat has been associated with a range of adrenal disorders, including APAs, where the majority of cases express high levels of βCat (*24*). To investigate whether high levels of βCat might contribute to increased AJ enrichment in human APAs, we analyzed paraffin-embedded sections from a cohort of 16 patients with APAs (**Supplementary Table 1**). We employed immunostaining for CYP11B2 (Aldosterone Synthase, AS), βCat, and KCad. Focusing first on AS-expressing regions, in a blinded fashion, we randomly assessed βCat expression levels and then independently scored AJ structures by determining the level of intercellular KCad (**Fig. 6A-B, Supplementary Fig. 9A**). While βCat expression levels showed no correlation with CYP11B2 expression in APAs, KCad peak intensity demonstrated a weak correlation (**Supplementary Fig. 9B-C**). Notably, APAs with higher βCat expression levels showed higher KCad levels at cell-cell contacts compared to those with lower βCat levels, which showed lower KCad levels (**Fig. 6A-C**). Using PLA as a proxy for AJ stability, we assessed the interaction between p120-KCad, which revealed ZA structures were enriched in APAs with elevated βCat levels (**Supplementary 9D**) mirroring the observations in βCat-GOF mice (**Fig. 4-5**). To quantitatively and spatially assess the AJs, we scored ∼50 cell pairs (across their cell-cell contacts) within each of 76 regions across all 16 APA samples for βCat levels (mean fluorescence intensity) and KCad levels (peak fluorescence intensity). Spearman correlation analysis comparing average data for each APA revealed a positive correlation between βCat levels and KCad levels (r=0.58) (**Fig. 6D**), with remarkable intratumor congruence (**Supplementary Fig. 9E**). Together, these data suggest that elevated βCat expression is associated with enhanced AJ formation and stability—characterized by increased KCad localization at cell-cell contacts, increased binding of p120 and KCad, and enrichment of ZA structures—supporting a role for βCat stabilization in promoting epithelial cohesion and junctional integrity in APAs. In addition, these data highlight AJs as potential targets for the development of new treatments for these tumors.

**Figure 6.**
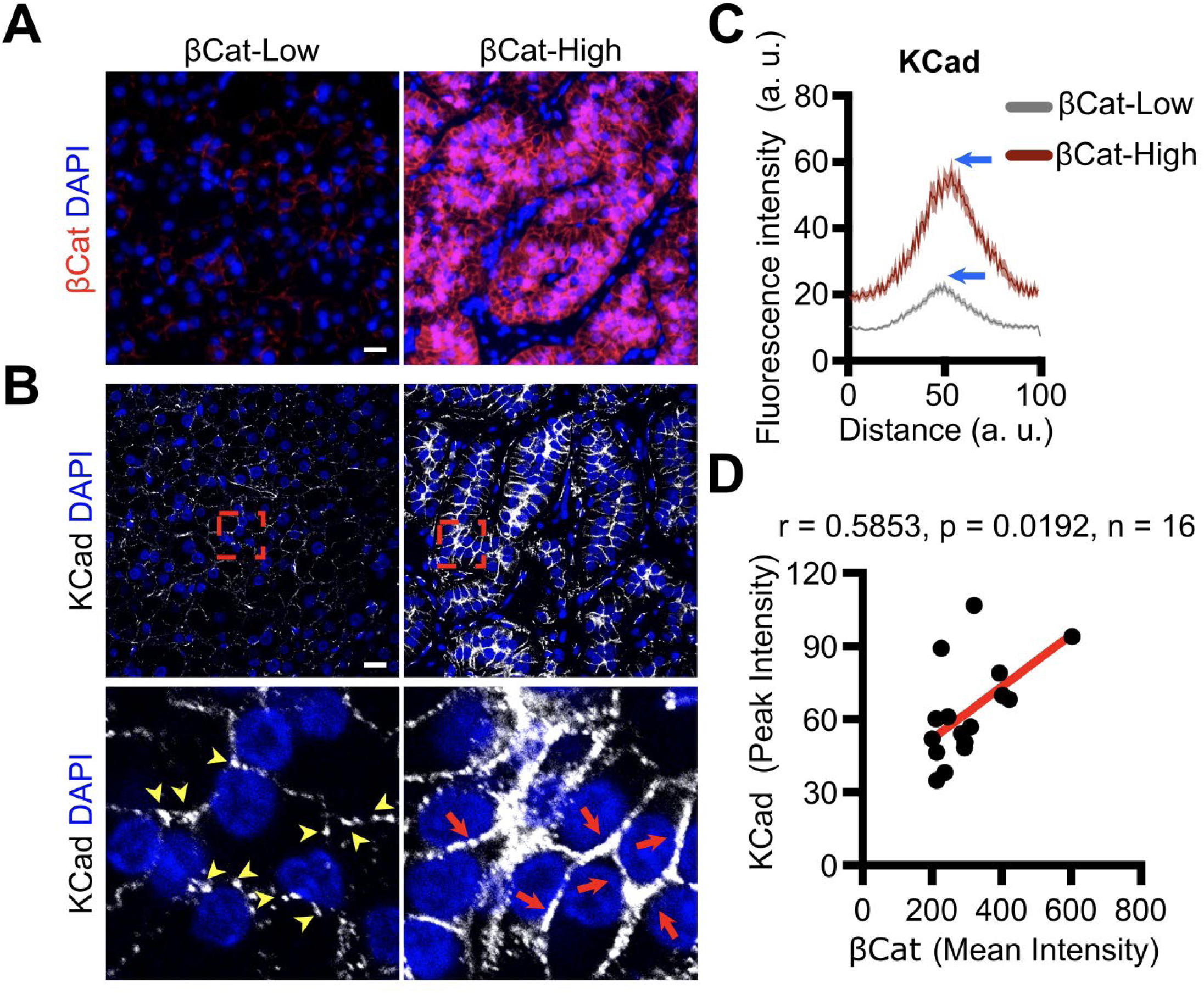
β-Catenin expression positively correlates with K-cadherin membrane localization in human aldosterone-producing adenomas. **A)** Representative images of β-catenin (βCat)-low and βCat-high immunofluorescence in human aldosterone-producing adenoma (APA) sections. **B)** Representative images of K-Cadherin (KCad) immunofluorescence from regions in (A), with magnified views of regions marked by dashed red squares. Yellow arrowheads denote puncta adherens junctions (pAJs), and red arrows denote zonula adherens (ZAs). **C)** Quantitative line profile analysis of KCad fluorescence intensity in representative βCat-low and βCat-high regions, with blue arrows highlighting peaks of averaged KCad signal localized at ∼50 cell-cell interfaces from randomly selected regions in APA sections. Lines represent mean and shaded areas represent standard error of the mean. **D)** Spearman correlation analysis comparing βCat mean intensity and peak KCad intensity, as shown in (C), from 16 human APA samples. The red line represents the regression fit (*r* = 0.5853, *P* < 0.0192). Scale bars, 20 μm.

## DISCUSSION

Aberrant aldosterone production underlies PA, a condition affecting millions worldwide with significant cardiovascular consequences. While the clinical importance of PA is well established, the molecular mechanisms governing zG expansion and hormone dysregulation remain incompletely understood. Our study addresses this knowledge gap by elucidating a novel signaling axis in the adrenal cortex that connects cell adhesion, tissue architecture, and hormone production. Specifically, we demonstrate that stabilization of βCat (βCat-GOF), beyond its recognized role as a transcription factor, drives AJ enrichment and subsequent zG morphogenesis (**Fig. 7**). Our findings reveal that zG architecture, particularly the specialized rosette structures unique to the zG, depends on AJ integrity for both morphological maintenance and coordinated hormone secretion. Under normal conditions, we have uncovered that physiological aldosterone secretagogues (K^+^ and AngII) activate the Rho-ROCK-NMII signaling pathway leading to enhanced AJ stability. In addition, employing models of βCat-GOF, we discovered how this pathway is hijacked under pathophysiological conditions to drive zG hyperplasia and excess hormone production. Moreover, using zG-specific mouse models and pharmacological inhibitors of ROCK signaling, we showed how this pathway can be targeted to mitigate zG hyperplasia and excessive aldosterone production. In contrast to other adrenal states that demonstrate sexual dimorphism (*26, 38–40*), no meaningful differences were noted between male and female mice in this study regarding their morphological response to fasudil, αCat-LOF, βCat-GOF or the combination of βCat-GOF and αCat-LOF. Together, these results underscore what appear to be fundamental regulatory mechanisms underlying adrenal homeostasis and the response to βCat-driven zG hyperplasia. Finally, the translational relevance of this work is underscored by our demonstration that βCat levels positively correlate with AJ formation in human APAs, further highlighting AJs as a potential therapeutic target for PA.

**Figure 7.**
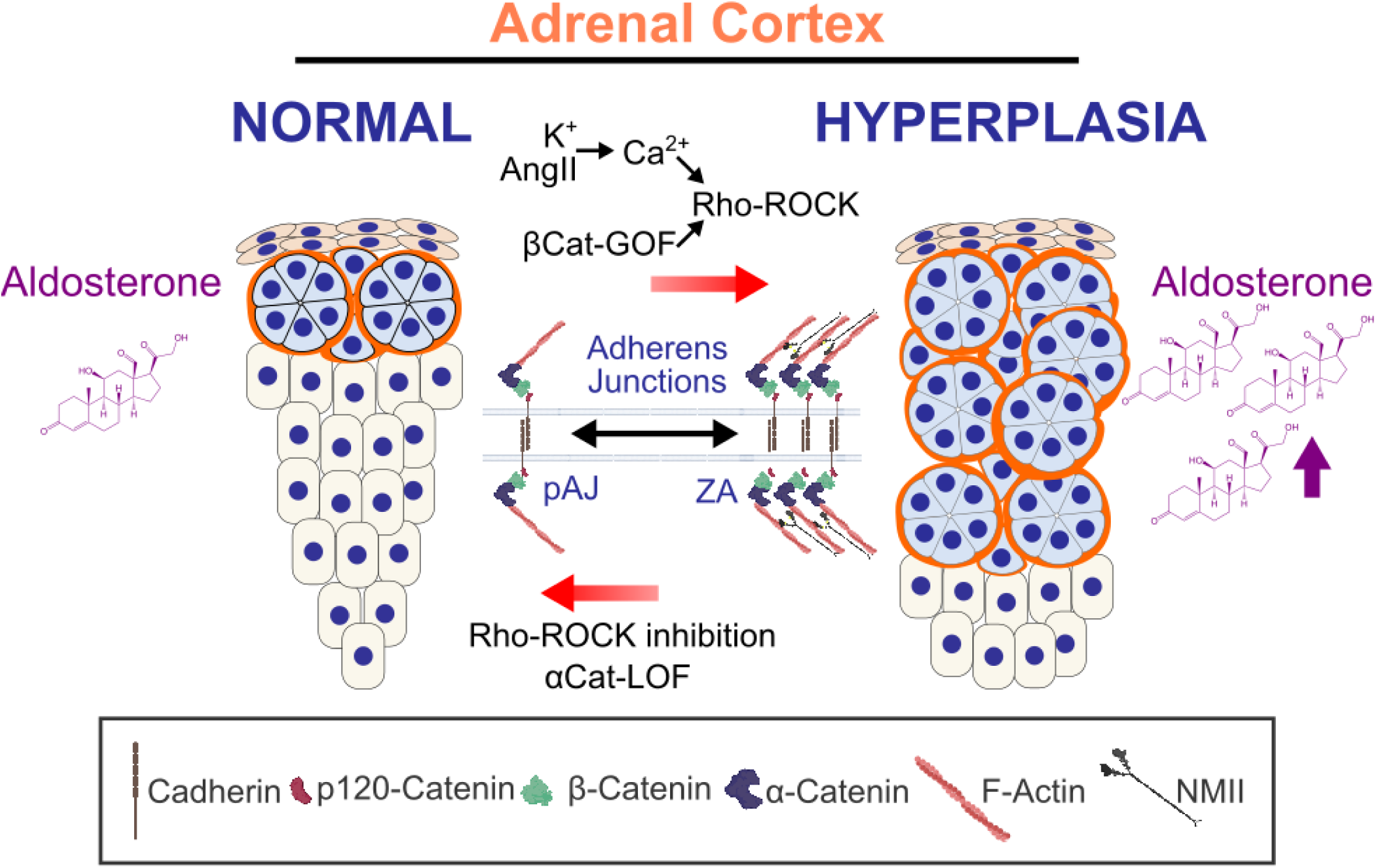
Schematic Illustrating Dynamic Regulation of Adherens Junctions in the Zona Glomerulosa. In the adrenal cortex, aldosterone secretagogues (K^+^ and AngII) and β-Catenin (βCat) gain-of-function (GOF) drives aldosterone production and hyperplasia of the zona Glomerulosa (zG) by stabilizing adherens junctions (AJs) through Rho-ROCK-actomyosin signaling, with α-Catenin (αCat) and Cadherins playing critical roles in this process. In mice, βCat-GOF increases AJ stability and aldosterone production, while Rho-ROCK inhibition or αCat deletion blunts these effects.

### βCat Stabilization Drives Adherens Junction Enrichment

Stabilization of βCat is reported in the majority of APAs (*24*), and mutations leading to βCat-GOF in mice drive zG hyperplasia (*41, 42*). βCat is localized in the nucleus, cytoplasm, and plasma membrane in zG cells (*24*) and is an essential component of the AJ. Here, we extend our prior analysis showing mouse adrenals with zG-specific βCat-GOF demonstrate upregulation of key genes that comprise the AJ, including *Cdh6* (KCad) and *Ctnnd1* (p120) (*10*). Now, we show direct interaction of p120 and KCad, a marker of a stable AJ, also increases following βCat-GOF in zG cells. The increase of KCad and p120 expression, along with an increase in their direct interaction following βCat-GOF, suggests that βCat regulates AJ stability both transcriptionally and through direct interactions at AJs. Moreover, treatment of the adrenocortical cell line NCI-H295R with CHIR, that leads to βCat stabilization, also increases membrane localization of AJ proteins, including KCad, αCat, and F-Actin, resulting in the formation of ZA. We further show that in human APAs βCat levels positively correlate with KCad levels and increased formation of ZAs. Although, these data are inherently correlative, it is reinforced by causal evidence from our cell-based and mouse genetic models showing that βCat stabilization directly enhances AJ stability and epithelial organization. This integrative approach supports the conclusion that βCat plays a functional role in shaping tumor architecture in APA via modulation of cell–cell adhesion. Together, these data indicate that βCat stabilization drives AJ enrichment and stability in mice and humans.

### Rho-ROCK Signaling Drives AJ Enrichment in the Adrenal

AJs are highly dynamic structures whose formation and stability are precisely regulated by Rho signaling and actomyosin contractility (*18, 43*). We have previously shown that AJs in the zG mediate intercellular communication via rosette structures to coordinate dynamic aldosterone release in response to physiological stimulation (*44*). But how such signals are transmitted from the extracellular space to the AJs remains poorly understood. Here, we extend our prior proteomic analysis of adrenal cells stimulated with K^+^, which identified increased levels of phosphoproteins regulated by Rho-GTPase activity (*26*). We demonstrate that stimulation of adrenal cells with K^+^ or AngII leads to the robust activation of the Rho-ROCK-NMII pathway and formation of ZAs via Ca^2+^ signaling at cell-cell contact points. Underscoring the role of this pathway, inhibition of Rho-ROCK (using Y27632 or fasudil) or NMII (using blebbistatin) effectively prevents actomyosin contraction and disrupts ZA formation following stimulation. These findings provide compelling evidence that physiological stimuli enhance AJ stability through activation of Rho-signaling, with additional kinases like myosin light chain kinase (MLCK) potentially contributing to this process. Consistent with this, human APAs have been identified with both increased abundance of RhoC (*45*) as well as missense mutations in Rho regulatory proteins (*46, 47*), supporting a model of autonomous pathological activation of this pathway. Collectively, our results establish a critical signaling axis through which aldosterone secretagogues trigger Rho activation in a physiological context, leading to enhanced AJ stability, cellular hyperplasia and regulation of aldosterone secretion—a mechanism that may be hijacked in APAs and in adrenal hyperplasia.

### Disruption of AJs impairs zG Rosettes and Aldosterone Production

During mammalian development, rosettes are essential for the morphogenesis of a range of tissues including the neural tube, kidney tubule, and pancreas, among others (*8, 48*). Rosettes primarily represent temporary developmental structures, which either resolve or open to form a lumen during mammalian development. Despite this, zG rosettes form during the first six weeks after birth in mice and are present throughout life (*10*). In addition, we have previously shown that zG rosettes function as the fundamental unit of the zG and coordinate calcium-mediated electrical activity evoked by the aldosterone secretagogue, AngII (*44*). Moreover, rosettes have also been described in the zG of adult humans, and larger rosettes have recently been associated with APMs (*10, 11*). To better understand the role of AJs in zG rosettes, we previously generated zG-specific βCat-LOF and βCat-GOF mice, which demonstrate marked disruption of rosettes in βCat-LOF adrenals and expansion of rosettes in βCat-GOF adrenals (*10*). However, βCat’s dual roles as a transcription factor and a structural component of the AJ (*20*) has limited our ability to fully delineate its impact on zG pathophysiology—particularly at the AJ. This duality also complicates efforts to isolate the specific contribution of AJs in this context. To address this challenge, we targeted αCat, a crucial AJ adaptor protein that lacks transcriptional activity, by generating zG-specific αCat-LOF mice. Consistent with the requirement for AJs in rosette formation, αCat-LOF mice demonstrate a decrease in zG rosette numbers without impacting the overall size of the zG. Similarly, inhibition of Rho-ROCK signaling (with fasudil), *in vivo,* also decreased the size of zG glomerular structures. Taken together, these findings indicate that AJs, Rho signaling, and αCat are essential for normal formation of zG rosette structures in adult mammals. In addition, these results underscore the role of zG morphology, and specifically the role of rosettes (rather than zG size *per se*), as being key to the level of aldosterone produced.

### βCat Drives Adrenal Hyperplasia via Enhanced AJ Formation

Constitutive activation of canonical Wnt/βCat signaling promotes tumorigenesis in multiple tissues, including the colon and the adrenal (*25, 49*). In addition, it is well known that nuclear βCat associates with TCF/LEF transcription factors to promote proliferation (*49, 50*). In contrast, we have previously shown that zG-specific βCat-GOF leads to zG hyperplasia without an increase in zG cell proliferation (*22*). To further explore this paradox, we assessed the role of AJs in zG-specific βCat-GOF mice using bigenic βCat-GOF::αCat-LOF mice. Consistent with our previous findings (*22*), βCat-GOF led to a progressive increase in zG hyperplasia. In contrast, morphological analysis of bigenic adrenals indicates that αCat, and by extension AJs, are required for βCat-driven zG hyperplasia. Interestingly, this requirement for αCat in βCat-driven hyperplasia mirrors findings in the Apc^Min^ mouse model of intestinal cancer, where simultaneous loss of αCat and Apc during tumor initiation suppresses adenoma formation(*51*), suggesting a conserved role for αCat in supporting early stages of βCat-mediated tissue expansion across different organ systems.

### AJ Disruption Sensitizes Hyperplastic zG Cells to Apoptosis

The increased cleaved caspase-3 immunostaining observed in both βCat-GOF::αCat-LOF adrenals and fasudil-treated βCat-GOF mice aligns with established mechanisms whereby AJ disruption influences cell survival. Previous studies have demonstrated that disruption of cadherin-catenin complexes can sensitize epithelial cells to apoptosis through multiple pathways, including exposing death receptors such as Fas to their ligands when AJ integrity is compromised(*35*), and through loss of the normal cellular microenvironment that supports cell survival(*37*). In the adrenal, zG cells are organized into specialized multicellular rosettes formed through AJ-mediated constriction. Disruption of this highly organized AJ-dependent architecture may be particularly effective at promoting apoptotic elimination of hyperplastic cells by destabilizing the structural integrity that normally supports cell survival, effectively counterbalancing the hyperplastic drive imposed by constitutive βCat activation. While these results indicate a role for AJ disruption in sensitizing cells to apoptosis, the complete molecular mechanisms linking junction destabilization to enhanced caspase activation in this specific rosette-organized tissue context warrant further investigation. Together, these studies highlight a critical role for AJs in the pathogenesis of βCat-driven zG hyperplasia in mice. Additional studies using mouse models of APAs will be essential to define the functional significance of AJs in adenoma formation.

### Molecular Heterogeneity and AJ Regulation in Human APAs

The βCat-GOF mouse model provides a valuable platform for investigating how zG hyperplasia contributes to PA. In humans, however, PA typically originates from somatic mutation–driven focal hyperplasia rather than from global βCat activation or diffuse hyperplasia. Notably, aberrant βCat stabilization occurs in roughly 70% of APAs, even though direct CTNNB1 mutations account for only 3–5% of cases. By contrast, somatic alterations in ion channels and pumps—particularly KCNJ5, CACNA1D, and ATP1A1—are far more prevalent, with KCNJ5 mutations alone comprising 38–70% of APAs across different populations(*52*). Interestingly, APAs harboring KCNJ5 mutations exhibit lower rates of βCat stabilization compared to other genotypes, highlighting distinct molecular pathways and suggesting potential subclassification within APAs. Moreover, some APAs—regardless of βCat status— harbor missense mutations in Rho GTPase–activating proteins(*47, 53*), which inactivate Rho signaling, or display upregulated RhoC expression, particularly in KCNJ5-mutant tumors(*45*). Collectively, these findings underscore the complex interplay of signaling networks in APA pathogenesis.

Correlation analyses revealed distinct relationships between these molecular markers in our cohort of 16 human APAs. While βCat expression levels showed no significant correlation with CYP11B2 expression, KCad peak intensity demonstrated a weak but detectable correlation with CYP11B2. Importantly, βCat expression levels strongly correlated with intercellular KCad levels. Although the sample size is limited in this analysis, this differential correlation pattern supports the interpretation that βCat’s primary role in APAs is independent of those mechanisms directly regulating aldosterone production. Instead, βCat likely contributes to adenoma development through its role in stabilizing AJs and promoting epithelial cohesion, consistent with the two-hit model of APA pathogenesis(*2, 23, 54*). The weak correlation between KCad and CYP11B2 may reflect their coordinated involvement in Ca^2+^-dependent signaling processes, as optimum aldosterone synthesis requires coordinated calcium bursts in response to secretagogues like AngII, and AJs facilitate intercellular Ca^2+^ coordination within the rosette architecture. Functionally, APAs with higher βCat expression levels showed significantly elevated KCad levels at cell-cell contacts compared to those with lower βCat levels. Furthermore, p120-catenin binding to KCad was enhanced in APAs with high βCat levels, indicating robust AJ stability in these tumors. p120-catenin plays a critical role in stabilizing cadherin-catenin complexes at the cell surface by preventing cadherin endocytosis and proteasomal degradation(*33*), thus providing a mechanistic basis for the observed correlation between βCat and AJ integrity. Collectively, current evidence highlights the critical involvement of AJ-associated pathways in human APA pathogenesis. However, the precise somatic mutation status of these tumor samples remains to be determined, warranting future studies leveraging expanded human APA cohorts with systematic mutational analyses.

### ROCK Inhibition Blocks zG Hyperplasia and Represents a Novel Therapeutic Approach for PA

ROCK inhibitors have demonstrated therapeutic potential across a wide range of conditions, including cardiovascular, neurodegenerative, ophthalmological, renal, metabolic, respiratory, and oncological disorders, among others (*55–60*). In addition, fasudil is well tolerated by humans and oral administration is currently being studied in a Phase 1 clinical trial (*61*). Despite this extensive effort, ROCK inhibitors have not been studied in the context of adrenal disorders, including PA, of which the vast majority of cases are caused by APMs or APAs. Current non-surgical therapeutic strategies for PA are focused on blocking excess aldosterone signaling or reducing aldosterone production (*62*). Our study demonstrates that Rho-ROCK inhibition (using fasudil) effectively attenuates zG hyperplasia and reduces aldosterone production. These findings indicate the zG is a direct target for ROCK inhibitors and suggest that ROCK inhibition may offer a promising new therapeutic strategy for treating PA. Finally, the known vasodilatory effects of these drugs may provide additional benefits to hypertensive patients with PA (*63*).

## METHODS

### Sex as a biological variable

Our study examined male and female animals, and similar findings are reported for both sexes. Please see details below and in the main text.

### Animals

Boston Children’s Hospital’s Institutional Animal Care and Use Committee approved all animal procedures. Mice were kept on a 12/12h light/dark cycle and had free access to standard chow and tap water.

Transgenic βCat-GOF (Cyp11b2^Cre/+^::Ctnnb1^flox(Ex3)/wt^), αCat-LOF (Cyp11b2^Cre/+^::Ctnna1^flox/flox^) and βCat-GOF::αCat-LOF (Cyp11b2^Cre/+^::Ctnnb1^flox(Ex3)/wt^::Ctnna1^flox/flox^) mice were generated by crossbreeding Cyp11b2^Cre/+^(*31*), Ctnna1^flox/flox^ (*64*) and Ctnnb1^flox(Ex3)/wt^ (*65*). Cyp11b2^Cre/+^::Ctnnb1^wt/wt^::Ctnna1^wt/wt^ mice were used as controls. Littermates were used whenever possible and efforts were made to avoid genetic drift between control and experimental strains. Mice between 6-12 weeks of age are classified as two months of age and mice between 16-21 weeks of age are classified as four months of age. The sex of the mice is specified in the corresponding figure legends.

### *In vivo* fasudil treatments

#### Young adult mice

Two-month-old C57BL6/J or littermate controls and βCat-GOF mice were injected intraperitoneally with 30 mg/kg fasudil (Selleckchem, dissolved in PBS) every other day for 28 days. Animals injected with PBS served as vehicle controls. The animals were sacrificed at the end of the experiments, and adrenals and kidneys were collected for immunofluorescence and mRNA expression analysis.

#### Adult mice

Four-month-old littermate βCat-GOF mice were injected intraperitoneally with 30 mg/kg fasudil or PBS six times per week for 28 days. For the last three days, mice were kept individually in metabolic cages for 24-hour urinary aldosterone measurements. The animals were sacrificed at the end of the experiments, and adrenals and kidneys were collected for immunofluorescence and mRNA expression analysis.

#### Aged mice

7-12-month-old littermate control and βCat-GOF mice were injected intraperitoneally with 30 mg/kg fasudil for 14 days. Three days before treatment and for the last three days of treatment, mice were kept individually in metabolic cages for 24-hour urinary aldosterone measurements.

### Urinary aldosterone measurements

Vehicle and fasudil-treated C57BL6/J and βCat-GOF mice were acclimated in metabolic cages for two days. Afterwards, their urine was collected for 24-hour periods using metabolic cages. Urinary aldosterone excretion was measured using a competitive radioimmunoassay Aldosterone kit (TECAN) following the manufacturer’s protocol. Briefly, 100 µl of urine was mixed with 1000 µl of hydrochloric acid and incubated at 30°C for 16 hours. Afterwards, 50 μl of this solution and 150 μl of the Zero Calibrator, together with 500 μl of the aldosterone radioactive tracer, were added to aldosterone antibody-coated tubes. After mixing, the tubes were incubated at room temperature for 18 hours. Then, the incubation mixture was decanted, and radioactivity in the tubes was counted using a Cobra II auto-gamma counter (Long Island Scientific, Berthold Technologies, New York, USA). Aldosterone levels were normalized to urinary creatinine measured using a calorimetric assay kit (Cayman Chemical).

### Cell culture

The NCI-H295R human adrenocortical carcinoma cell line (passage 6-15) was maintained in DMEM-F12 (GIBCO) supplemented with 2.5% Nu-Serum (Fisher Scientific), 1X insulin-transferrin-selenium (ITS) (GIBCO), 1X Glutamate, and 1X penicillin/streptomycin.

### Immunofluorescence assays

Antibodies used in these studies are listed in **Supplementary Table 2.**

#### Cell culture

NCI-H295R cells were seeded onto poly-L-lysine (Sigma)-coated coverslips placed in 24-well plates. Following one day in culture, cells were serum-starved in DMEM/F12 medium containing 1% ITS for 16 hours. Cells were then treated for 1 hour with either 10 µM (–) blebbistatin (StemCell Technologies), 50 µM Y27632 (Tocris), 10 µM fasudil (Selleckchem), 10 µM nifedipine (Sigma), 20 µM iCRT14 (MedChemExpress) or DMSO as a control. Following pretreatment, cells were stimulated with either 11 mM KCl, 100 nM AngII, 5 µM CHIR 99021, or vehicle controls (11 mM NaCl for KCl and DMSO for the small molecules). AngII was diluted in an 11 mM NaCl solution, allowing NaCl to serve as the vehicle control for both KCl and AngII treatments.

After 48 hours of stimulation (unless otherwise specified), cells were fixed with 4% paraformaldehyde (PFA) for 15 minutes at room temperature and washed three times with PBS. Cells were permeabilized with PBS-0.1%Tween (PBS-T) for 15 minutes and blocked with 1% BSA in PBS-T for 20 minutes. Samples were incubated with primary antibodies (**Supplementary Table 2)** diluted in blocking buffer for 4 hours at room temperature. After three PBS washes, cells were incubated for 2 hours with fluorophore-conjugated secondary antibodies (1:400 dilution) (**Supplementary Table 2)**, far-red fluorescent phalloidin conjugate (1:1000; Life Technologies), CellMask Actin Tracking stain (Invitrogen), and 4′,6-diamidino-2-phenylindole (DAPI; 1:1000; Sigma). Finally, coverslips were washed three times with PBS and mounted onto glass slides using ProLong Gold antifade mountant (Invitrogen).

#### Thick sections

For imaging of rosette structures, adrenal glands were collected from mice and immediately placed in cold PBS. After carefully removing the surrounding fat tissue, the glands were cut in half with a blade and fixed overnight at 4°C in 4% PFA. The fixed adrenals were then embedded in 4% low-melting temperature agarose, and 70-μm-thick sections were prepared using a Leica Vibratome and transferred to 48-well plates. Tissue sections were permeabilized with PBS-T for 30 minutes and then blocked with a solution of 1% BSA and 5% normal goat serum (Sigma) in PBS-T for one hour. Samples were incubated with primary antibodies diluted in blocking buffer overnight. After three washes with PBS-T, sections were incubated for two hours with fluorophore-conjugated secondary antibodies (1:400 dilution) and DAPI. Following three additional washes with PBS-T, adrenal slices were mounted on slides using ProLong Gold antifade mountant (Invitrogen).

#### Paraffin sections

Immunofluorescent analysis was performed using established protocols, as described previously(*10*). In brief, after fixation, adrenals were dehydrated in ethanol and embedded in paraffin blocks. Paraffin sections were cut at 5 µm thickness. Sections were rinsed in xylene, an ethanol gradient and then PBS. Antigen retrieval was performed in Tris-EDTA pH 9.0. Sections were blocked in 5% Normal Goat Serum, 0.1% Tween-20 in PBS for 1 h at RT. Primary antibodies were diluted 1:200 in 5% NGS in PBS and incubated on sections at 4 °C overnight. Slides were washed three times for 5 min in 0.1% Tween-20 in PBS. Secondary antibodies were diluted in 1:200 in PBS and incubated on sections at RT for 1–2 h. For nuclear staining, DAPI (4′,6-diamidino-2-phenylindole) was added to the secondary antibody mixture at a final concentration of 1:1000. After three 5-min washes with 0.1% Tween-20 in PBS, slides were mounted with ProLong Gold Antifade Mountant (Thermo Fisher Scientific, P36930). Antibodies are listed in Supplemental Table 2.

### Proximity Ligation Assay

After deparaffinization and antigen retrieval, adrenal paraffin sections were permeabilized with PBS-T for five minutes and blocked with normal goat serum (NGS) for one hour. Afterward, the sections were incubated with KCad and p120 antibodies (1:200 in NGS), overnight at 4°C. Proximity ligation assays were performed using Anti-Rabbit PLUS, Anti-Mouse MINUS PLUS probes combined with the FarRed Detection Reagents kit (DuoLink) following the manufacturer’s protocol. Incubation with no primary antibody, only KCad primary antibody and only p120 primary antibody were used as negative controls.

### Image acquisition and quantification

Imaging was performed using a Zeiss Axio Imager Z2, a Zeiss LSM 510, or a Zeiss LSM 700 Laser Scanning Confocal microscope.

PLA signal intensity in the zG area, determined by the density of nuclei, was calculated after subtracting background using ImageJ software (National Institute of Health) and the measure tool.

Equatorial sections of adrenal glands were imaged for analysis when possible encompassing the capsule, cortical regions, and the medulla. Analysis of the Dab2(+) area was used as a proxy for the zG and was normalized to the whole cortical area using Zeiss Zen 3.9 software and presented as Dab2(+) Area (% Cortex). The two-dimensional area of glomerular structures, identified by closed circular Laminin β1 labeling, was manually quantified using Zeiss ZEN 3.9 image analysis software (ZEISS). Rosettes were identified as Laminin β1-encapsulated glomerular structures containing ≥5 DAPI-stained nuclei.

The numbers of Ki67(+) and Cleaved-Cas3(+) cells within the Dab2(+) area were manually counted and then normalized to the total Dab2(+) area (mm^2^), providing a density of positive cells per unit area.

### Quantification of βCat and KCad signal in human APA samples

Aldosterone-producing adenoma (APA) samples from patients diagnosed with primary aldosteronism were used in this study **(Supplementary Table 1)**. Consecutive paraffin sections of human APA tissue were subjected to immunofluorescence staining for CYP11B2, βCat, and KCad. Whole-slide images of immunostaining for CYP11B2 and βCat were acquired using a Zeiss Axio Imager Z2 microscope. For each sample, random CYP11B2(+) regions (measuring 255 x 255 µm) were selected in a double-blinded manner to minimize selection bias. Within these regions, the mean βCat and CYP11B2 fluorescence intensity was quantified using the Zen 3.9 analysis tools (ZEISS). In parallel, images of KCad immunostaining were acquired from the same regions using a Zeiss LSM 700 Laser Scanning Confocal microscope under standardized acquisition settings. KCad fluorescence intensity at cell-cell contact points was measured using the line profile tool in ImageJ, with quantification performed in a blinded fashion (as detailed below).

### Quantitative line profile analysis

The fluorescence signal intensity was measured along a line (50 pixels wide) drawn between two cell nuclei using the line profile tool in ImageJ. Raw data were organized in a tabular format, with each measurement consisting of paired columns representing height and signal values. To facilitate downstream analysis, a custom R function was developed to systematically extract every two adjacent columns from the data frame and convert each pair into a separate data frame. This ensured that each pair of height and signal measurements was handled independently for subsequent normalization and analysis.

To enable comparison across datasets with varying height ranges, all height measurements were linearly interpolated to a common scale ranging from 0 to 100 using a custom R function. For each paired data frame, the function identified the minimum and maximum height values and applied a linear transformation to rescale the heights to the standardized interval. Signal values were retained without modification. Data points containing missing values in either the height or signal columns were excluded from further analysis to ensure data integrity.

Following normalization, the list of processed data frames was combined into a single data frame using a custom R function. The resulting data frame contained all normalized height and signal data, structured for efficient batch analysis. To summarize the signal response across the normalized height range, the combined data were partitioned into 1-unit intervals from 0 to 100. The mean signal value within each interval for every measurement pair was computed. For each interval, the function selected all signal values whose corresponding normalized heights fell within the interval and calculated the average, excluding missing values.

### Fluorescence Recovery After Photobleaching

Viral particles were prepared using pLV-Puro-EF1A-hCDH6-3xGGGGS-GFP transfer plasmid containing the CDH6-GFP fusion construct along with packaging plasmids pLP1, pLP2, and pVSVG in HEK293T cells. H295R cells were transduced with lentiviral particles carrying the pLenti-KCad-GFP plasmid in the presence of 5 μg/mL polybrene. Cells were incubated with viral particles for 72 hours. KCad-GFP expressing H295R cells were seeded onto 35 mm glass-bottom dishes.

Two days prior to FRAP experiments, media was changed to serum-starved media overnight. Cells were then treated with either NaCl or KCl (15 mM final concentration) and maintained in serum-starved conditions until experiments. One hour prior to the experiment, media was changed to imaging buffer containing (in mM): HEPES 5, NaCl 120, Na2HPO4 1.6, NaH2PO4 5.4, CaCl2 1.3, MgCl2 1, KCl 3, glucose 15, pH 7.4. For K^+^ stimulated cells, NaCl in the buffer was exchanged with KCl 15 mM (final concentration) to maintain osmolarity. Using a Leica TCS SP8 Laser Scanning Confocal with a 63× oil immersion objective, KCad-GFP expressing cells were monitored using the FRAP wizard. The FRAP protocol consisted of a pre-bleaching phase for 50 seconds with 5-second intervals (10 frames); a bleaching phase, during which a region of the cell-cell junction was bleached for 1.3 seconds, repeated twice, and a recovery phase for 10 minutes with 5-second intervals (120 frames). Fluorescence recovery after photobleaching was analyzed by normalizing the recovered intensity in the bleached ROI relative to the pre-bleaching and immediate post-bleach mean intensities. The normalization formula used was:

F(t)normalized = 100 × [F(t) - F0] / [Fpre - F0]

where F(t) is the fluorescence intensity at time t, F0 is the immediate post-bleach intensity, and Fpre is the pre-bleach intensity. Recovery curves were fitted using nonlinear regression in GraphPad Prism 10 to determine the mobile fraction and recovery half-time (t1/2).

### RNA isolation and qPCR

NCI-H295R cells were plated in 12-well plates. Following one day in culture, cells were then pretreated for one hour with either iCRT14 (MedChemExpress) or DMSO as the control. Following pretreatment, cells were stimulated with either 5 µM CHIR 99021 or vehicle (DMSO) for 48 hours. Whole tissue (adrenal, kidney) was homogenized with a tissue homogenizer, and total RNA and cDNA were prepared as described previously (*26*). Gene expression analysis was assessed by TaqMan Universal PCR Master Mix (Applied Biosystems). Taqman probes are listed in **Supplementary Table 3.**

### Coimmunoprecipitation

Cells were seeded in 100 mm culture dishes at 80-90% confluency and serum-starved for 16 hours in DMEM/F12 medium supplemented with 1× ITS. Following starvation, cells were stimulated with KCl (11 mM), AngII (100 nM) or NaCl (11 mM) (vehicle) for 48 hours. After stimulation, cells were lysed on ice using IP lysis buffer supplemented with protease and phosphatase inhibitors (Santa Cruz Biotechnology). Lysates were clarified by centrifugation at 16,000 × *g* for 10 minutes at 4°C, and the supernatant was collected.

For immunoprecipitation, 50 μL of Protein A/G beads were washed and equilibrated three times with IP lysis buffer. Subsequently, 100 μL of cell lysate and 10 μL of anti-αCat antibody were added to the beads and incubated overnight at 4°C with gentle rotation. The beads were then washed three times with IP lysis buffer. Immunoprecipitated proteins were eluted in 50 μL of 2× Laemmli buffer and samples were heated at 95°C for 15 minutes. Samples were stored at −20°C until Western blot analysis.

### Active Rho-GTP pull-down assay

Cells were seeded in 100 mm culture dishes and serum-starved for 16 hours in DMEM/F12 medium supplemented with 1× ITS. Following starvation, cells were stimulated with KCl (11 mM), AngII (100 nM) or NaCl (11 mM) (vehicle) for three hours. Rho-GTP was pulled down with the RhoA Pull-Down Activation Assay Biochem Kit (Cytoskeleton), following the manufacturer’s protocol.

### Western blot

Protein lysates prepared in Laemmli buffer were boiled at 95°C for 10 minutes, separated by sodium dodecyl sulfate-polyacrylamide gel electrophoresis (SDS-PAGE), and transferred to polyvinylidene difluoride (PVDF) membranes. Membranes were blocked with either 1% bovine serum albumin (BSA) or 5% skim milk powder in PBS-T for 15 minutes, then incubated with primary antibodies overnight at 4°C. After three washes with PBS-T, membranes were incubated with horseradish peroxidase (HRP)- conjugated secondary antibodies in blocking solution for 1 hour at room temperature. Following three additional PBS-T washes, membranes were developed using enhanced chemiluminescence (ECL) substrate. Chemiluminescent signals were detected either by exposure to X-ray film or by imaging with a ChemiDoc Imaging System (Bio-Rad). Densitometric analysis of band intensities was performed using ImageJ software.

### Gene Enrichment Analysis

Significantly deregulated phosphoproteins (FoldChange > 2, FDR < 0.1) following K⁺ stimulation of NCI-H295R cells were analyzed for molecular function Gene Ontology (GO) terms and Reactome pathway enrichment using EnrichR(*66*).

### Statistical Analysis

No power analyses were conducted for this study. All statistical analyses were performed using Prism 10 software (GraphPad). No data were excluded from the analysis. Specific details regarding statistical tests and sample sizes (n), which represent independent mice, samples, or experiments, are provided in the corresponding figure legends. Data are presented as mean ± SEM unless otherwise indicated.

### Study approval

All animal procedures were approved by Boston Children’s Hospital’s Institutional Animal Care and Use Committee. The studies done with human adrenal samples were approved by the ethics commission of the Canton of Zurich, Switzerland (BASEC-Nr. 2017-00771). After obtaining written informed consent, we collected adrenal tissues from patients undergoing adrenalectomy for aldosterone-producing adenomas at the University Hospital Zurich. The diagnosis of PA/aldosterone-producing adenomas was established in accordance with institutional and international guidelines.

## Acknowledgement

We thank members of the Breault laboratory, the Division of Endocrinology at Boston Children’s Hospital and Oguz C. Koc for helpful comments and suggestions and Dr. Mark Taketo for the gift of Ctnnb1fl(ex3) mice. This work was supported by 2R01DK123694 (to DTB), the Swiss National Science Foundation (310030L_182700/1) (to FB) and by the University Research Priority Program of the University of Zurich (URPP) ITINERARE – Innovative Therapies in Rare Diseases (to FB).

