## Supplementary Figures for "Rho-ROCK signaling and α-Catenin mediate β-Catenin-driven hyperplasia in the adrenal via adherens junctions"

**Supplementary Figure 1**

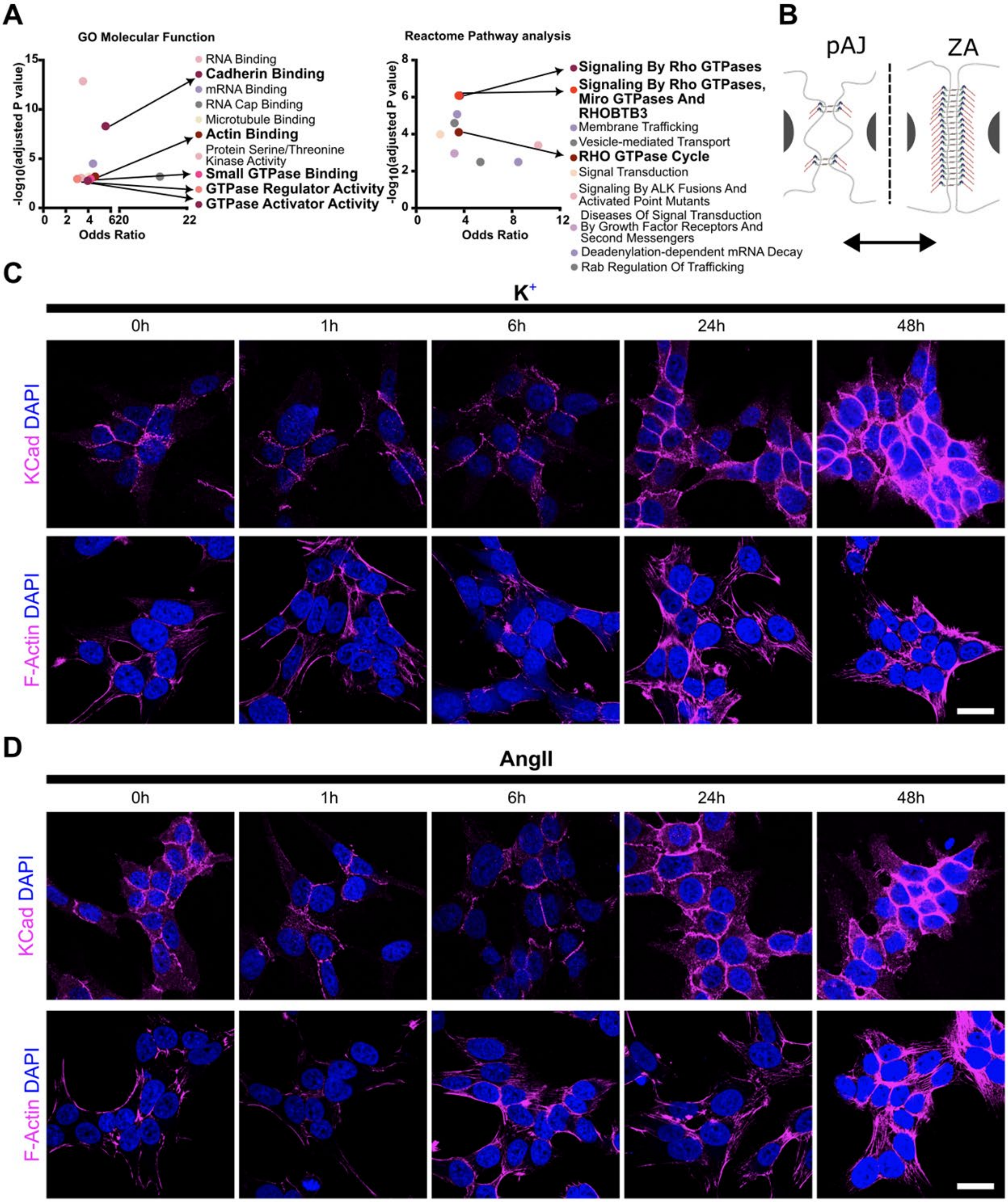

**Supplementary Figure 1: Rho signaling pathways and adherens junctions are regulated upon secretagogue stimulation.**

**A)** Significantly enriched molecular function gene ontology (GO) terms and Reactome pathways of dysregulated phosphoproteins upon $K^+$ stimulation in NCI-H295R cells. **B)** Schematic of puncta adherens junctions (pAJ) and zonula adherens (ZA) in epithelial cells. **C)** Time-course analysis of K-Cadherin (KCad) and filamentous actin (F-Actin) immunofluorescence upon $K^+$ stimulation (0-48 hr) in NCI-H295R cells. **D)** Time-course analysis of KCad immunofluorescence and F-Actin staining upon AngII stimulation (0-48 hr) in NCI-H295R cells. Representative images are shown. Scale bar, 20 μm.

**Supplementary Figure 2**

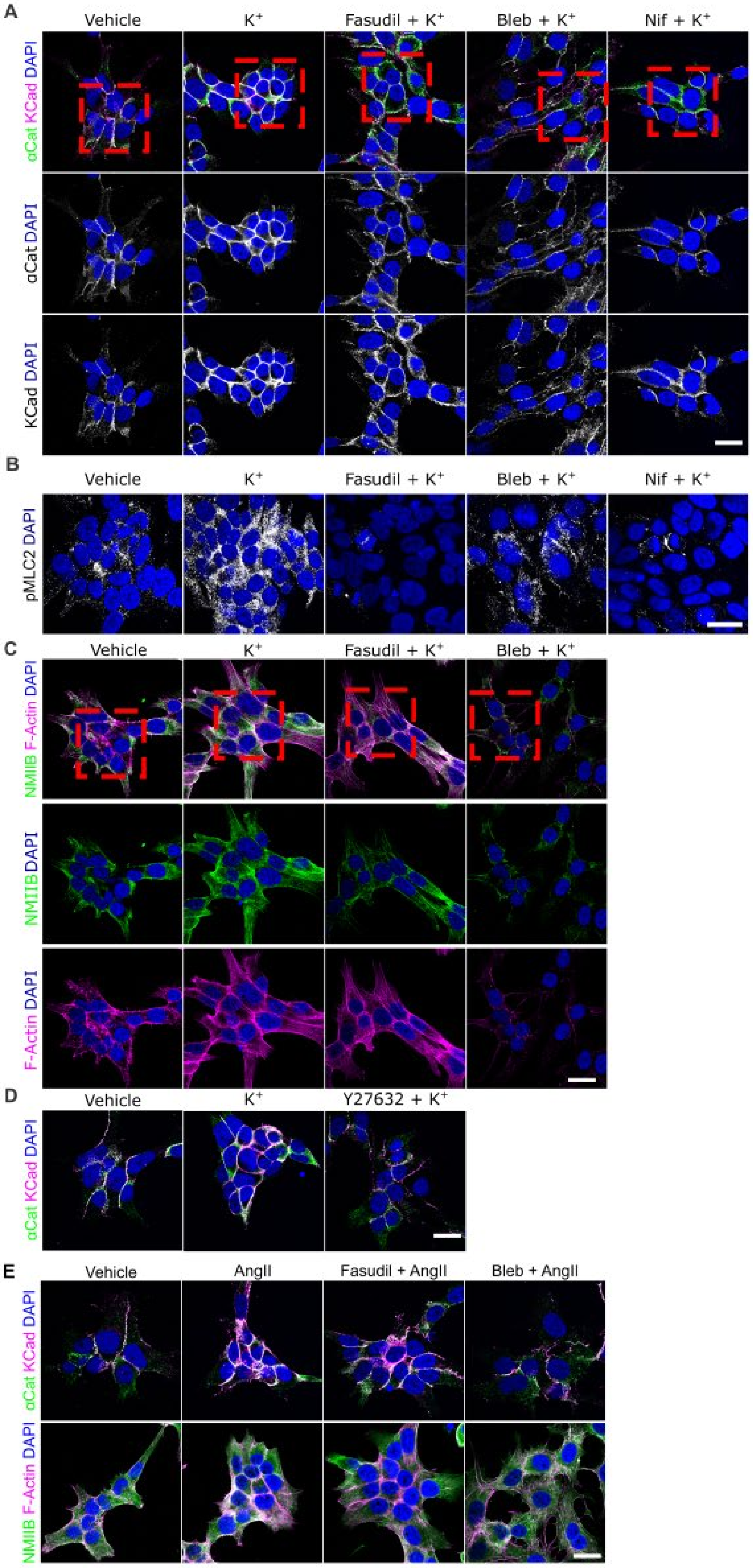

**Supplementary Figure 2: Aldosterone secretagogues increase adherens junction stability in NCI-H295R cells.**

**A)** Representative images of α-Catenin (αCat) and K-Cadherin (KCad) immunofluorescence in NCI-H295R cells stimulated with vehicle (11 mM NaCl) and potassium ($K^+$;11 mM KCl) ± fasudil (10 µM), blebbistatin (10 µM) or nifedipine (10 µM) for 48 h (1 h preincubation with inhibitors). Red dashed squares demarcate regions magnified in Figure 1D. **B)** Representative images of phospho-Myosin Light Chain2 (pMLC2) immunofluorescence in NCI-H295R cells treated as in (A). **C)** Representative images of NMIIB immunofluorescence and F-Actin staining in NCI-H295R cells treated as in (A). Red dashed squares demarcate regions magnified in Figure 1H. **D)** Representative images of αCat and KCad immunofluorescence in NCI-H295R cells stimulated with vehicle (11 mM NaCl), $K^+$ (11 mM) ± Y27632 (50 µM). **E)** Representative images of αCat, KCad, non-muscle myosin IIB (NMIIB) immunofluorescence and filamentous actin (F-Actin) staining in NCI-H295R cells stimulated with vehicle (11 mM NaCl), AngII (100 nM) ± fasudil (10 µM) or blebbistatin (10 µM) for 48 h (1 h preincubation with inhibitors). Scale bars, 20 µm.

**Supplementary Figure 3**

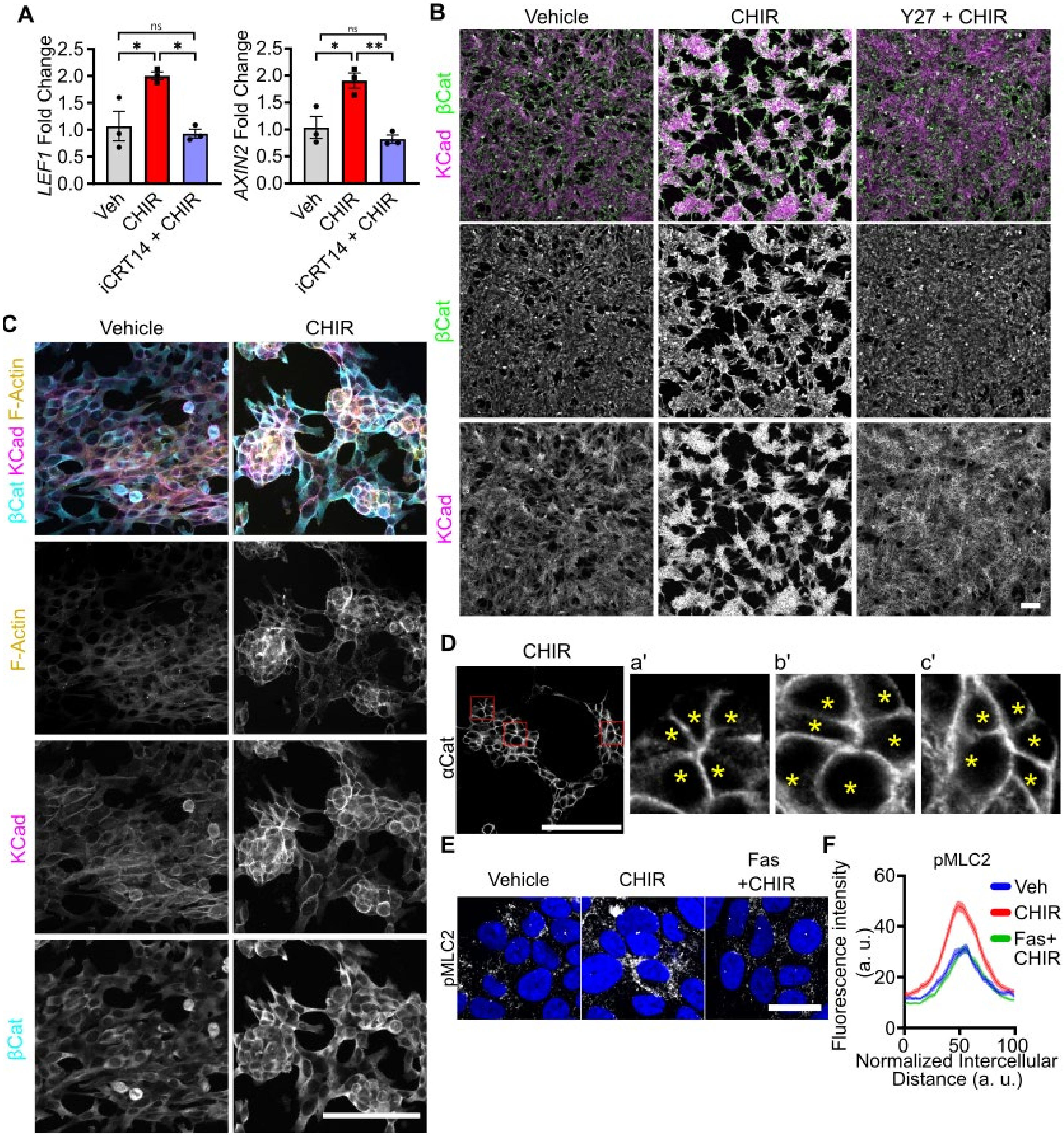

**Supplementary Figure 3: β-Catenin Stabilization via CHIR stimulation leads to NCI-H295R cell aggregation and formation of rosette-like structures.**

**A)** *LEF1* and *AXIN2* mRNA expression in H295R with CHIR 99021 (CHIR; 5 µM) ± iCRT14 (20 µM) or vehicle for 48 h (1 h preincubation with iCRT14). Statistical significance determined by one-way ANOVA with Tukey's multiple-comparison posttest (*P < 0.05, **P < 0.01, ns, not significant). Data are presented as mean ± SEM. **B)** Representative images of β-Catenin (βCat) and K-Cadherin (KCad) immunofluorescence in NCI-H295R cells stimulated with CHIR ± Y27632 (Y27) or vehicle (DMSO) for 48 h (1 h preincubation with Y27). **C)** Representative Z-stack projections showing βCat, KCad immunofluorescence and filamentous-actin (F-Actin) localization in NCI-H295R cells treated with CHIR (5 µM) or vehicle. **D)** Representative images of α-Catenin (αCat) immunofluorescence in NCI-H295R cells stimulated with CHIR. Areas highlighted by red squares are enlarged in panels a', b', and c'. Each asterisk denotes an individual cell within an NCI-H295R rosette-like structure. **E)** Representative images of phospho-Myosin Light Chain2 (pMLC2) immunofluorescence in NCI-H295R cells with CHIR ± Fasudil (Fas) or vehicle (DMSO) for 48 h (1 h preincubation with Fas). F) Quantitative line profile analysis pMLC2 fluorescence intensities represented in (E). Lines represent mean and shaded areas represent standard error of the mean (SEM). Scale bars, 100 µm or 20 µm (in E).

**Supplementary Figure 4**

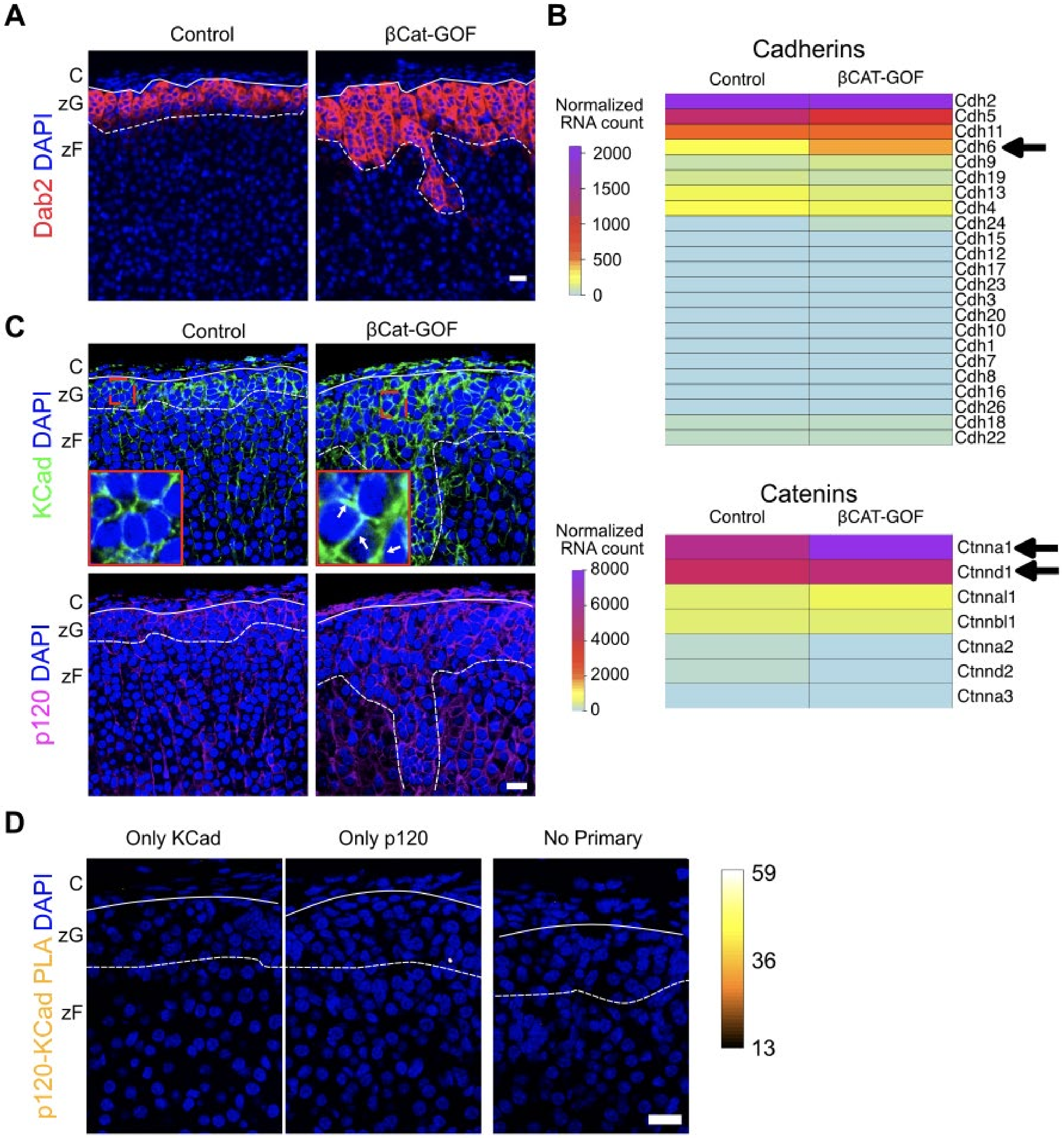

**Supplementary Figure 4: Adherens junction proteins are upregulated in βCat-GOF mice.**

**A)** Representative images of Dab2 immunofluorescence in adrenal sections from two-month-old control and βCat-GOF female mice. **B)** Expression profile of cadherin and catenin protein family in the adrenal glands of control and βCat-GOF mice (GEO: GSE144503). Arrows indicate upregulated genes in βCat-GOF adrenals. **C)** Representative images of KCad and p120-Cat (p120) immunofluorescence in adrenal sections from control and βCat-GOF female mice, with magnified views of regions marked by dashed red squares. The arrows denote the zonula adherens. **D)** Representative images of proximity ligation assay negative controls. The fluorescence signal is represented as a heat map using an orange-hot lookup table. Solid lines mark the boundary between the adrenal capsule (C) and zona Glomerulosa (zG), while dashed lines indicate the boundary between the zG and zona fasciculata (zF). Scale bars, 20 μM.

**Supplementary Figure 5**

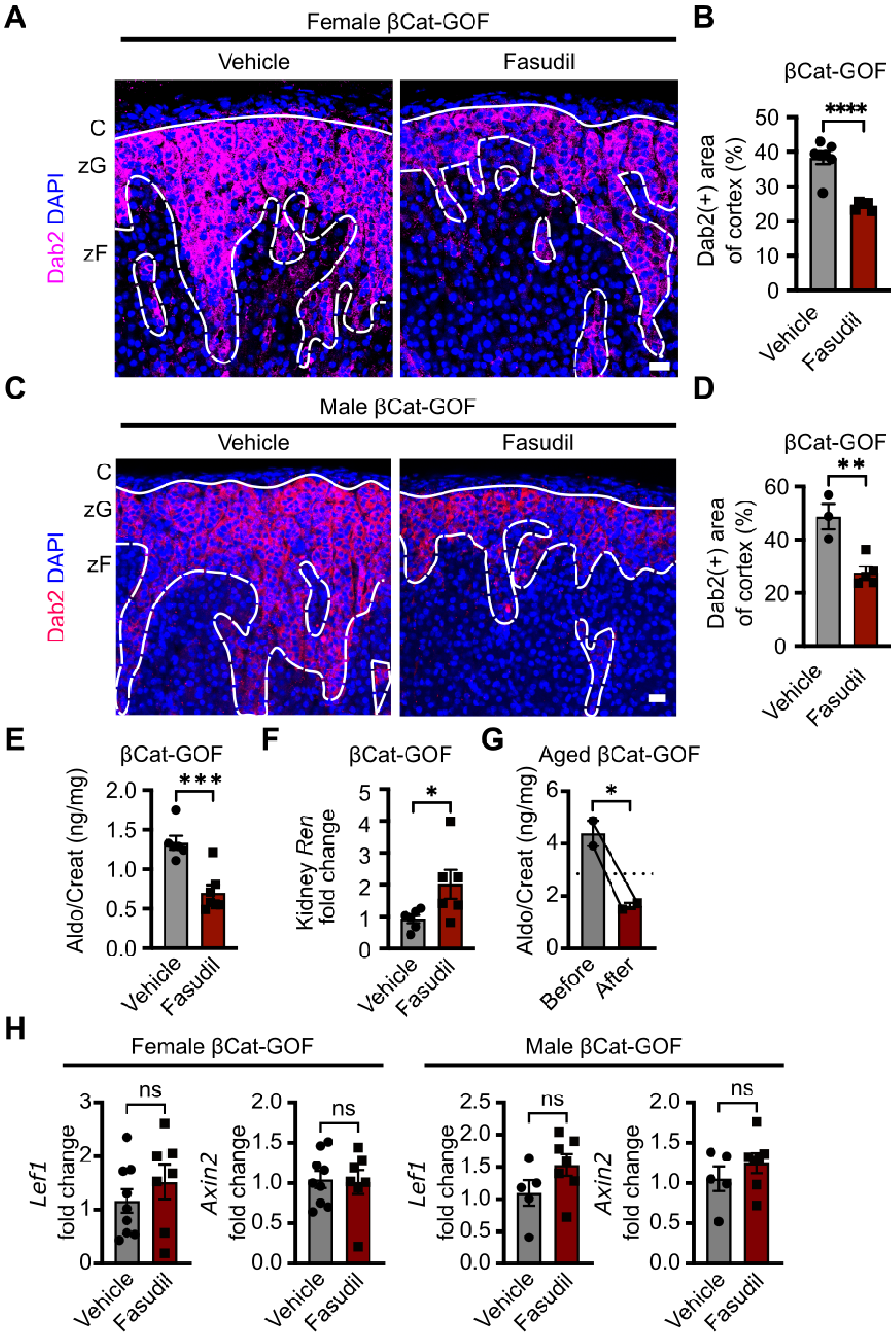

**Supplementary Figure 5: ROCK inhibition with fasudil prevents hyperplasia and reduces aldosterone production.**

**A)** Representative images of Dab2 immunolabeling in adrenal sections from vehicle- or fasudil-treated four-month-old βCat-GOF female mice. **B)** Quantification of the percentage of Dab2(+) area in the adrenal cortex, as shown in (A) (n = 7, 5 mice). **C)** Representative images of Dab2 immunolabeling in adrenal sections from vehicle- or fasudil-treated four-month-old βCat-GOF male mice. **D)** Quantification of the percentage of Dab2-positive area in the adrenal cortex, as shown in (A) (n = 3, 5 mice). **E)** Measurement of 24-hour urinary aldosterone levels in vehicle- or fasudil-treated four-month-old βCat-GOF male mice, assessed by radioimmunoassay (RIA) and normalized to creatinine (n = 6, 7 mice). **F)** Renin mRNA expression in kidneys of vehicle- and fasudil-treated four-month-old βCat-GOF male mice, assessed by quantitative PCR (qPCR) (n = 6, 6 mice). **G)** Measurement of 24-hour urinary aldosterone levels in one-year-old βCat-GOF male mice before and after fasudil treatment, assessed by RIA and normalized to creatinine (n = 2 mice). Dotted line indicates mean aldosterone levels in untreated littermate controls. Data points for controls are presented in Figure 2H. **H)** *Lef1* and *Axin2* mRNA expression in kidneys of vehicle- and fasudil-treated four-month-old βCat-GOF female and male mice, assessed by qPCR (n = 5, 9 mice). Statistical significance determined by unpaired two-tailed Student's *t*-test or ratio paired t-test (in G) (*P < 0.05, **P < 0.01, ***P < 0.001, ****P < 0.0001). Data are presented as mean ± SEM. Solid lines mark the boundary between the adrenal capsule (C) and zona glomerulosa (zG), while dashed lines indicate the boundary between the zG and zona fasciculata (zF). Scale bars, 20 μm.

**Supplementary Figure 6**

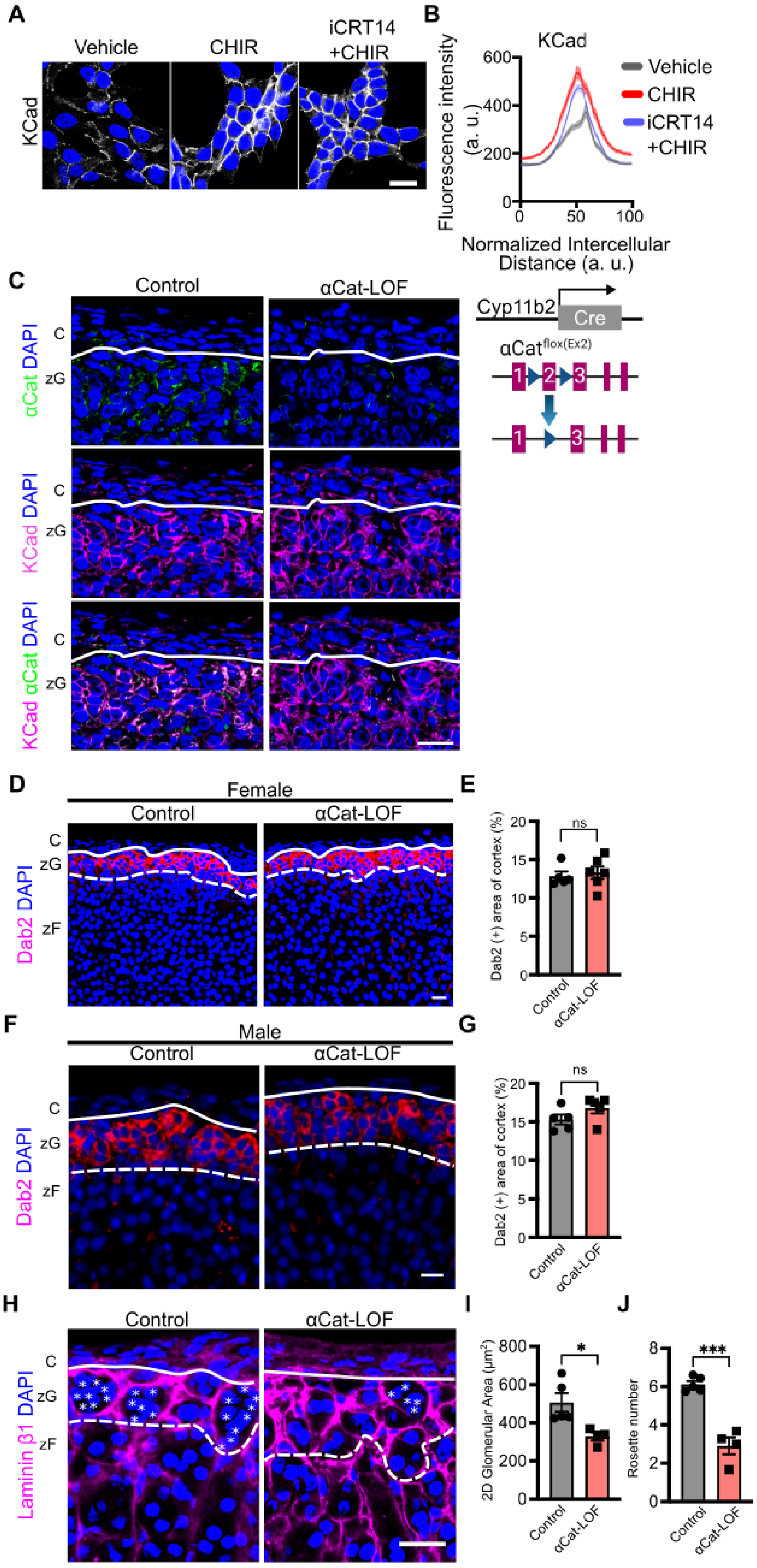

**Supplementary Figure 6: zG-specific αCat deletion reduces rosette numbers.**

**A)** Representative images of K-Cadherin (KCad) immunofluorescence in NCI-H295R cells with CHIR ± iCRT14 (20 µM) or vehicle (DMSO) for 48 h (1 h preincubation with iCRT14). **B)** Quantitative line profile analysis KCad fluorescence intensities represented in (A). Lines represent mean and shaded areas represent standard error of the mean (SEM). **C)** Representative images of αCat and KCad immunofluorescence of adrenal sections from two-month-old control and αCat-LOF male mice. Schematic of Cyp11b2$^{Cre/+}$::Ctnna1$^{flox/flox}$ (αCat-LOF) mice. **D)** Representative images of Dab2 immunolabeling in adrenal sections from two-month-old control and αCat-LOF female mice. **E)** Quantification of Dab2(+) percentage of the adrenal cortex represented in (B) (n = 5, 6 mice). **F)** Representative images of Dab2 immunolabeling in adrenal sections from two-month-old control and αCat-LOF male mice. **G)** Quantification of Dab2(+) percentage of the adrenal cortex represented in (D) (n = 5, 5 mice). **(H)** Representative images of Laminin β1 immunolabeling in adrenal sections from two-month-old control and αCat-LOF male mice. Each white asterisk denotes an individual cell within a zG rosette. **I)** Quantification of the two-dimensional (2D) area of glomerular structures, as defined by Laminin β1 labeling in (F) (n = 5, 4 mice). **J)** Quantification of rosette number, defined as clusters of five or more cells within a single glomerular structure in regions measuring 200 x 200 µm, as shown in (F). Each asterisk denotes an individual cell within a zG rosette (n = 5, 4 mice). All statistical significance determined by unpaired two-tailed Student's *t*-test (*P < 0.05, *** < 0.001, ns, not significant). Data are presented as mean ± SEM. Solid lines mark the boundary between the adrenal capsule (C) and zona glomerulosa (zG), while dashed lines indicate the boundary between the zG and zona fasciculata (zF). Scale bars, 20 µM.

**Supplementary Figure 7**

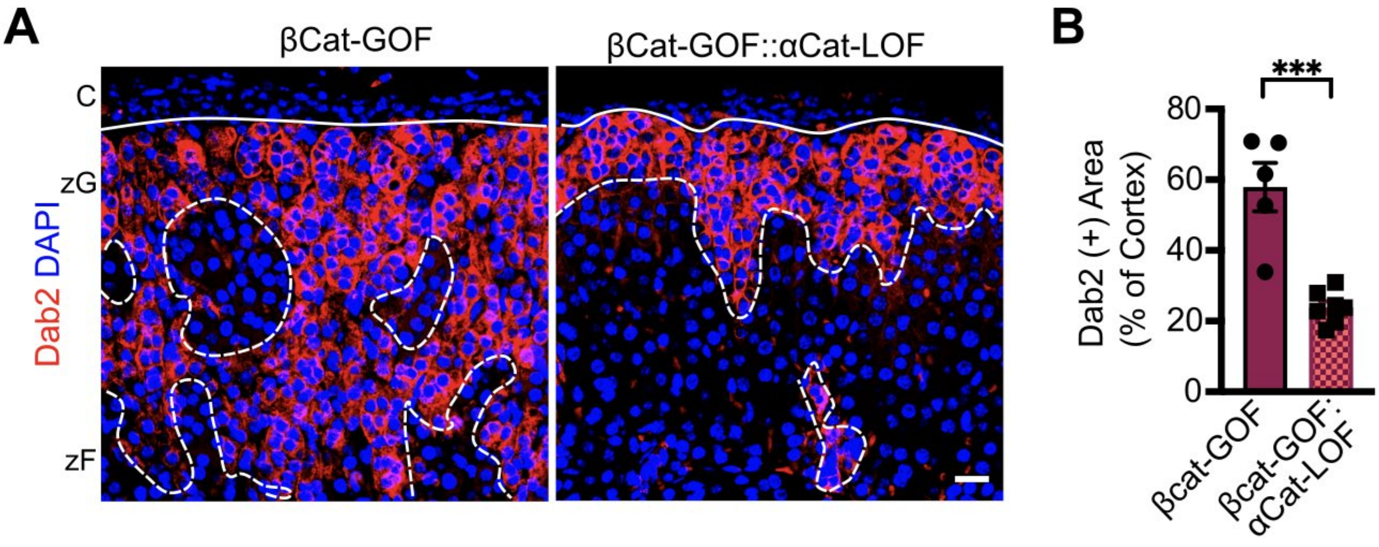


**Supplementary Figure 7: zG-specific α-Catenin deletion attenuates βCat-GOF-induced zG hyperplasia.**

**A)** Representative images of Dab2 immunolabeling in adrenal sections from four-month-old βCat-GOF and βCat-GOF::αCat-LOF female mice. **B)** Quantification of the percentage of Dab2(+) area in the adrenal cortex, represented in (A) (n = 5, 7 mice). Statistical significance determined by unpaired two-tailed Student's *t*-test (***P < 0.001). Data are presented as mean ± SEM. Solid lines mark the boundary between the adrenal capsule (C) and zona glomerulosa (zG), while dashed lines indicate the boundary between the zG and zona fasciculata (zF). Scale bar, 20 μM.

### Supplementary Figure 8

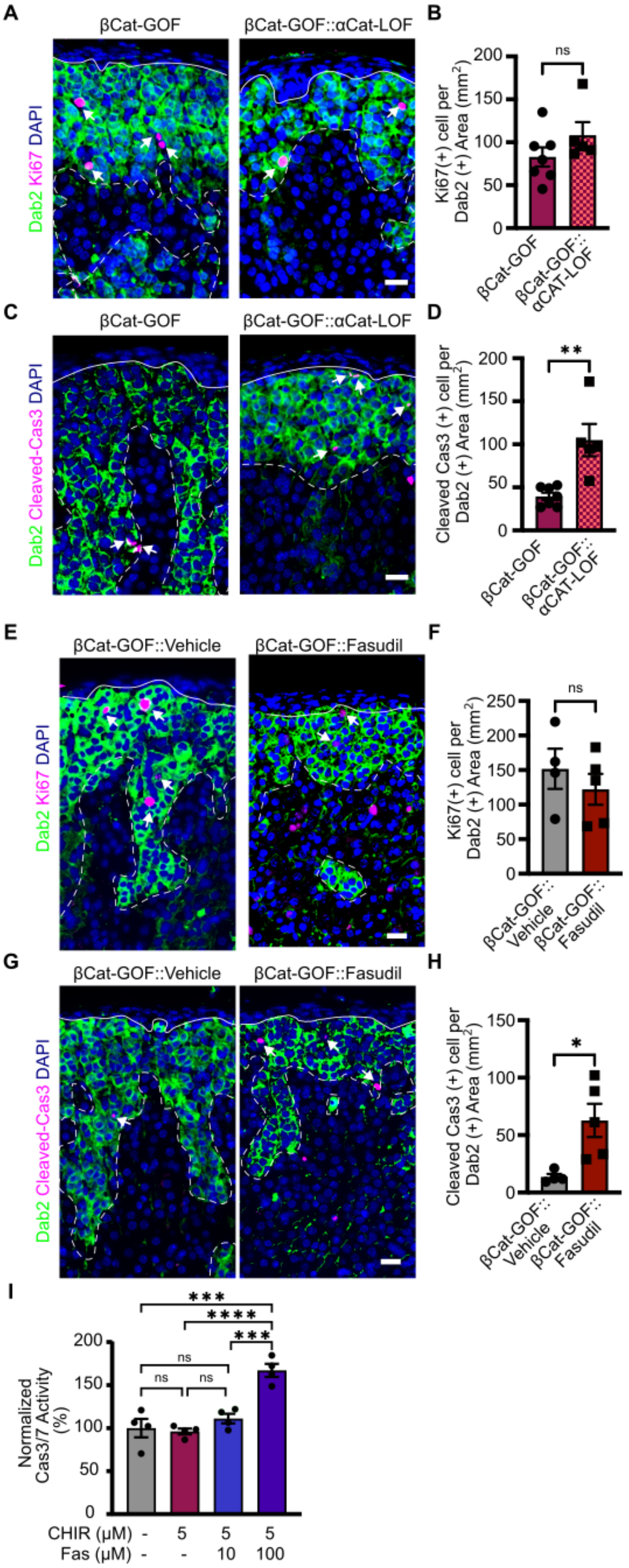

**Supplementary Figure 8: α-Catenin protects against apoptosis in βCat-GOF adrenals.**

**A)** Representative images of Dab2 and Ki67 immunolabeling in adrenal sections from four-month-old βCat-GOF and βCat-GOF::αCat-LOF male mice. **B)** Quantification of Ki67(+) cells in the Dab2(+) area, represented in (A) (n = 7, 5 mice). **C)** Representative images of Dab2 and cleaved-Caspase 3 immunolabeling in adrenal sections from four-month-old βCat-GOF and βCat-GOF::αCat-LOF male mice. **D)** Quantification of cleaved-Caspase 3-positive cells in Dab2(+) area, represented in (C) (n = 7, 5 mice). **E)** Representative images of Dab2 and Ki67 immunolabeling in adrenal sections from four-month-old βCat-GOF male mice treated with either vehicle (PBS) or fasudil (30 mg/kg) administered six times per week for 28 days. **F)** Quantification of Ki67(+) cells in the Dab2(+) area, represented in (E) (n = 4, 5 mice). **G)** Representative images of Dab2 and cleaved-Caspase 3 immunolabeling in adrenal sections from four-month-old βCat-GOF male mice treated as in (E). **H)** Quantification of cleaved-Caspase 3(+) cells in the Dab2(+) area, represented in (G) (n = 4, 5 mice). **I)** Caspase 3/7 luminescence activity measured in NCI-295R cells treated with CHIR (5μM) ± fasudil (Fas; 10 and 100 μM) or vehicle (DMSO) for 48 h (1 h preincubation with Fas) using Caspase-Glo 3/7 Assay System. Statistical significance determined by unpaired two-tailed Student's *t*-test or one-way ANOVA with Tukey's multiple-comparison posttest (in I) (*P < 0.05, **P < 0.01, ***P < 0.001, ****P < 0.0001, ns, not significant). Solid lines mark the boundary between the adrenal capsule (C) and zona glomerulosa (zG), while dashed lines indicate the boundary between the zG and zona fasciculata (zF). Scale bars, 20 μM.

**Supplementary Figure 9**

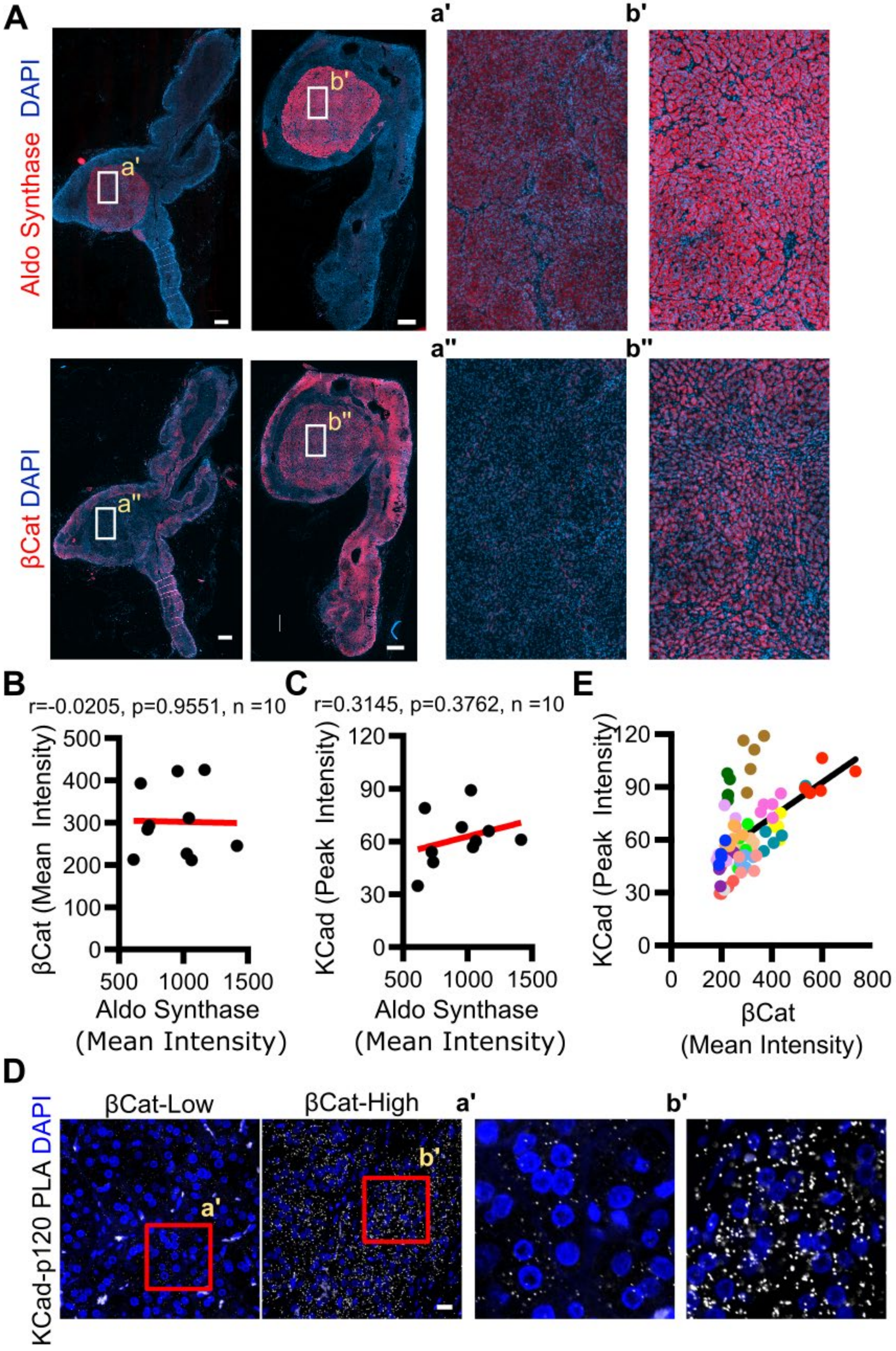

**Supplementary Figure 9: Correlation of β-Catenin and K-Cadherin expression in human aldosterone-producing adenomas.**

**A)** Representative immunofluorescence images showing aldosterone (Aldo) synthase and β-Catenin (βCat) labeling in human aldosterone-producing adenoma specimens from Figure 6. Areas highlighted by white rectangles are shown at higher magnification in panels a', b', a” and b''. Scale bars, 1 mm. **B-C)** Pearson correlation analysis comparing βCat mean intensity and Aldo synthase mean intensity **(B)** and K-Cadherin (KCad) mean intensity and Aldo synthase mean intensity **(C)** from 10 human APA samples. The red lines represent the regression fit. **D)** Representative images of KCad/p120 proximity ligation assay (PLA) signal in human aldosterone-producing adenoma sections. Areas highlighted by red rectangles are shown at higher magnification in panels a' and b'. Scale bar, 20 μM. **E)** Scatter plot showing quantification from each randomly selected region (n=3-5) for every tumor section, with different colors representing different individuals, from the analysis in Figure 6D comparing βCat mean intensity and KCad peak intensity.

**Supplementary Table 1: Information regarding patients harboring aldosterone-producing adenomas (APAs) assessed for β-Catenin and K-Cadherin correlation analysis.**

| Patient # | Sex | Age | Diagnosis |
|---|---|---|---|
| 1 | Female | 30 | Primary Aldosteronism (APA) |
| 2 | Female | 38 | Primary Aldosteronism (APA) |
| 3 | Male | 40 | Primary Aldosteronism (APA) |
| 4 | Male | 42 | Primary Aldosteronism (APA) |
| 5 | Male | 49 | Primary Aldosteronism (APA) |
| 6 | Female | 49 | Primary Aldosteronism (APA) |
| 7 | Male | 50 | Primary Aldosteronism (APA) |
| 8 | Male | 51 | Primary Aldosteronism (APA) |
| 9 | Male | 51 | Primary Aldosteronism (APA) |
| 10 | Male | 52 | Primary Aldosteronism (APA) |
| 11 | Female | 54 | Primary Aldosteronism (APA) |
| 12 | Male | 56 | Primary Aldosteronism (APA) |
| 13 | Male | 57 | Primary Aldosteronism (APA) |
| 14 | Female | 61 | Primary Aldosteronism (APA) |
| 15 | Male | 64 | Primary Aldosteronism (APA) |
| 16 | Male | 68 | Primary Aldosteronism (APA) |

**Supplementary Table 2: Antibodies list.**

| Antibody | Host | Source | Catalog # | Used for |
|---|---|---|---|---|
| anti-α-Catenin | Mouse | Invitrogen | 13-9700 | IP, IF, WB |
| anti-K-Cadherin | Rabbit | Abcam | ab133632 | IF, WB, PLA |
| anti-non-muscle-myosin II-B | Rabbit | BioLegend | 909901 | IF |
| anti-Laminin β1 | Rat | Santa Cruz | sc-33709 | IF |
| anti-β-Catenin | Mouse | BD Biosciences | 610154 | IF |
| anti-p120-Catenin | Mouse | BD Biosciences (Fisher Scientific) | BDB610133 | IF, PLA |
| anti-p120-Catenin | Mouse | Assay Biotech | YM4837 | PLA (Human) |
| anti-Dab2 | Rabbit | Cell Signaling Technologies | 12906 | IF |
| anti-Dab2 | Mouse | BD Biosciences | 610464 | IF |
| anti-Ki67 | Rabbit | Cell Signaling Technologies | 12202 | IF |
| anti-cleaved-Caspase3 | Rabbit | Cell Signaling Technologies | 9661 | IF |
| anti-Cyp11b2 | Mouse | Dr. Celso E. Gomez-Sanchez, University of Mississippi Medical Center, MS, USA | hCYP11B2-41-13B | IF |
| anti-β-Actin | Mouse | Santa Cruz | sc-47778 | WB |
| anti-Rabbit IgG-647 | Goat | Invitrogen | A21245 | IF |
| anti-Rabbit IgG-488 | Goat | Invitrogen | A11008 | IF |
| anti-Mouse IgG-488 | Donkey | Invitrogen | A21202 | IF |
| anti-Mouse IgG-647 | Donkey | Invitrogen | A31571 | IF |
| anti-Rat IgG-647 | Goat | Invitrogen | A21247 | IF |
| anti-Rabbit IgG-HRP | Goat | Cell Signaling Technologies | 7074 | WB |
| anti-Mouse IgG-HRP | Horse | Cell Signaling Technologies | 7076 | WB |

**Supplementary Table 3: TaqMan gene expression assay list.**

| Species | Gene | Assay catalog number |
|---|---|---|
| Human | *18S* | Hs99999901_s1 |
| Human | *AXIN2* | Hs00610344_m1 |
| Human | *LEF1* | Hs01547250_m1 |
| Mouse | *Gapdh* | Mm99999915_g1 |
| Mouse | *18s* | Mm02601777_g1 |
| Mouse | *Axin2* | Mm00443610_m1 |
| Mouse | *Lef1* | Mm00550265_m1 |
| Mouse | *Renin* | Mm02342887_mH |
| Mouse | *Cyp11b2* | Mm01204955_g1 |
